# NuRD independent Mi-2 activity represses ectopic gene expression during neuronal maturation

**DOI:** 10.1101/2022.04.19.488642

**Authors:** Gabriel N Aughey, Elhana Forsberg, Krista Grimes, Shen Zhang, Tony D Southall

## Abstract

During neuronal development, extensive changes to chromatin states occur to regulate lineage-specific gene expression. The molecular factors underlying the repression of non-neuronal genes in differentiated neurons are poorly characterised. The Mi2/NuRD complex is a multiprotein complex with nucleosome remodelling and histone deacetylase activity. Whilst NuRD has previously been implicated in the development of nervous system tissues the precise nature of the gene expression programmes that it coordinates are ill-defined. Furthermore, evidence from several species suggests that Mi-2 may be incorporated into multiple complexes that may not possess histone deacetylase activity. We show that Mi-2 activity is required for suppressing ectopic expression of germline genes in neurons independently of HDAC1/NuRD, whilst components of NuRD, including Mi-2, regulate neural gene expression to ensure proper development of the larval nervous system. We find that Mi-2 binding in the genome is dynamic during neuronal maturation and Mi-2 mediated repression of ectopic gene expression is restricted to the early stages of neuronal development, indicating that Mi-2/NuRD is required for establishing stable neuronal transcriptomes during the early stages of neuronal differentiation.

## Introduction

Neuronal development involves the transition of multipotent precursor cells into highly specialised neurons with distinct morphological and biochemical characteristics. Extensive gene expression changes accompany the differentiation of precursors into specialised cell-types. These gene expression changes are underpinned by a dynamic chromatin environment in which regulation of nucleosome positioning and histone modifications ensures appropriate transcriptional responses. Whilst recent studies have provided illuminating descriptions of changes to chromatin accessibility and histone modifications during the process of neuronal differentiation (Aughey et al., 2018; Janssens et al., 2022; Preissl et al., 2018; Ziffra et al., 2021), the chromatin-interacting complexes that affect these changes are not well defined.

The Mi-2/NuRD complex is a multiprotein chromatin remodelling complex that influences gene expression via two separate enzymatic activities; ATP-dependent nucleosome remodelling, and histone deacetylation (Tong et al., 1998; Wade et al., 1998; Xue et al., 1998; Zhang et al., 1998). These activities are mediated by enzymes found in two distinct subunits. The nucleosome remodelling subunit, Mi-2, is a well conserved ATP-dependent nucleosome remodeller (CHD4 in mammals). While histone deacetylation is conferred by a separate HDAC (histone deacetylase) enzyme. NuRD components are widely conserved and broadly expressed in most metazoan tissues, reflecting the important role of the complex in gene regulation. NuRD has been reported to have critical roles in developmental processes, (e.g, in muscle (Gomez-Del Arco et al., 2016) and hematopoietic (Yoshida et al., 2008) cell differentiation), as well as regulation of pluripotent cell reprogramming (Rais et al., 2013).

Mi-2/NuRD complexes have critical roles in neurodevelopment and have been implicated in the aetiology of neurodevelopmental disorders (Hoffmann and Spengler, 2019). A recent study showed that Mi-2 is required in *Drosophila* neuronal lineages to deactivate notch responsive stem-cell enhancers in neuronal progeny, thereby ensuring the fidelity of differentiated cell fate in neurons (Zacharioudaki et al., 2019). Similarly, NuRD was shown to deactivate gene expression by acting on promoters of presynaptic genes to promote synaptic connectivity (Yamada et al., 2014). These studies suggest a predominantly suppressive role for NuRD in the regulation of gene expression. Whilst this is consistent with early models of NuRD activity as a repressive complex, more recent studies have presented a more nuanced role for NuRD in modulating gene expression, which may be neither activating or repressive, but acting to fine tune gene expression and reduce transcriptional noise (Bornelov et al., 2018; Burgold et al., 2019; Ragheb et al., 2020). Therefore, an important unanswered question is whether NuRD is primarily repressive in neuronal development, or whether it may facilitate control of active gene expression as demonstrated in stem cell models.

NuRD composition is dynamic in mouse cortical development, with alternate Mi-2/CHD paralogues incorporated to mediate distinct effects in different developmental stages (Nitarska et al., 2016). In *Drosophila* the components of the NuRD complex are well conserved, (although only single homologues of most NuRD proteins are identified). However, despite the strong conservation of NuRD constituent proteins, and ample evidence for physical interactions between them (Reddy et al., 2010; Zhang et al., 2016), the extent to which the canonical NuRD complex regulates gene expression in *Drosophila* is uncertain. A separate complex consisting of only Mi2 and MEP-1 (termed dMec), has been suggested to be more prevalent than NuRD in fly, and may be responsible for the majority of Mi-2 dependent phenotypes (Kunert et al., 2009). This observation has not been corroborated by further studies, however, NuRD-independent roles for Mi-2 may be present in human cells (Amaya et al., 2013). Alternatively, it is possible that since NuRD is comprised of pre-assembled modules (Zhang et al., 2016), and Mi-2 may exist as a peripheral component to the core complex (Low et al., 2016), the detection of non-NuRD associated Mi-2 may simply be a proteomic artefact that does not reflect a *bona fide* biological activity. Therefore, the functional role of dMEC, and its relationship to the canonical NuRD holocomplex, remains poorly characterised.

Here, we set out to better understand the influence of Mi-2/NuRD on gene expression in the development of the central nervous system. Genomic profiling with Targeted DamID revealed distinct binding of Mi-2 containing complexes, NuRD and dMEC. Knockdown of Mi-2 in larval neurons resulted in ectopic expression non-neuronal genes as well as defective optic lobe development. Mi-2 represses lineage-inappropriate gene expression independently of NuRD, whilst optic lobe phenotypes are phenocopied by other members of the NuRD complex. We find that Mi-2 is not required for repression of non-neuronal genes after the early stages of differentiation, indicating that Mi-2 activity is critical for ensuring maintenance of the neuronal gene expression programme during maturation, but alternative mechanisms may be employed in fully differentiated neurons.

## Results

### *Mi-2* is required in neurons for survival and larval locomotion

*Mi-2* has been shown to protect neural lineages from mitogenic notch signalling, indicating that NuRD has an important role in gene regulation in the neuroblasts and early progeny of *Drosophila* central nervous system (CNS) lineages. To better understand NuRD function in the fly CNS, we set out to examine the distribution of the core ATP-dependent nucleosome remodelling subunit Mi-2. We utilised a MiMIC insertion line in which Mi-2 is EGFP-tagged at its endogenous locus *(Mi-2-GFP* hereafter). We observed widespread Mi-2 expression with detectable EGFP signal in multiple cell-types in the third instar CNS (Figure 1A). EGFP signal was visible in nuclei of neural stem cells (NSCs) (as distinguished by their larger size and absence of Elav or Repo) as well as all neurons and some glia, (Figure 1A). The nuclear expression of Mi-2 in progenitor and post-mitotic cell-types in the larval CNS suggests that Mi-2 activity is likely to be involved in gene regulation in all stages of neuronal differentiation, including differentiated neurons.

**Figure 1.**
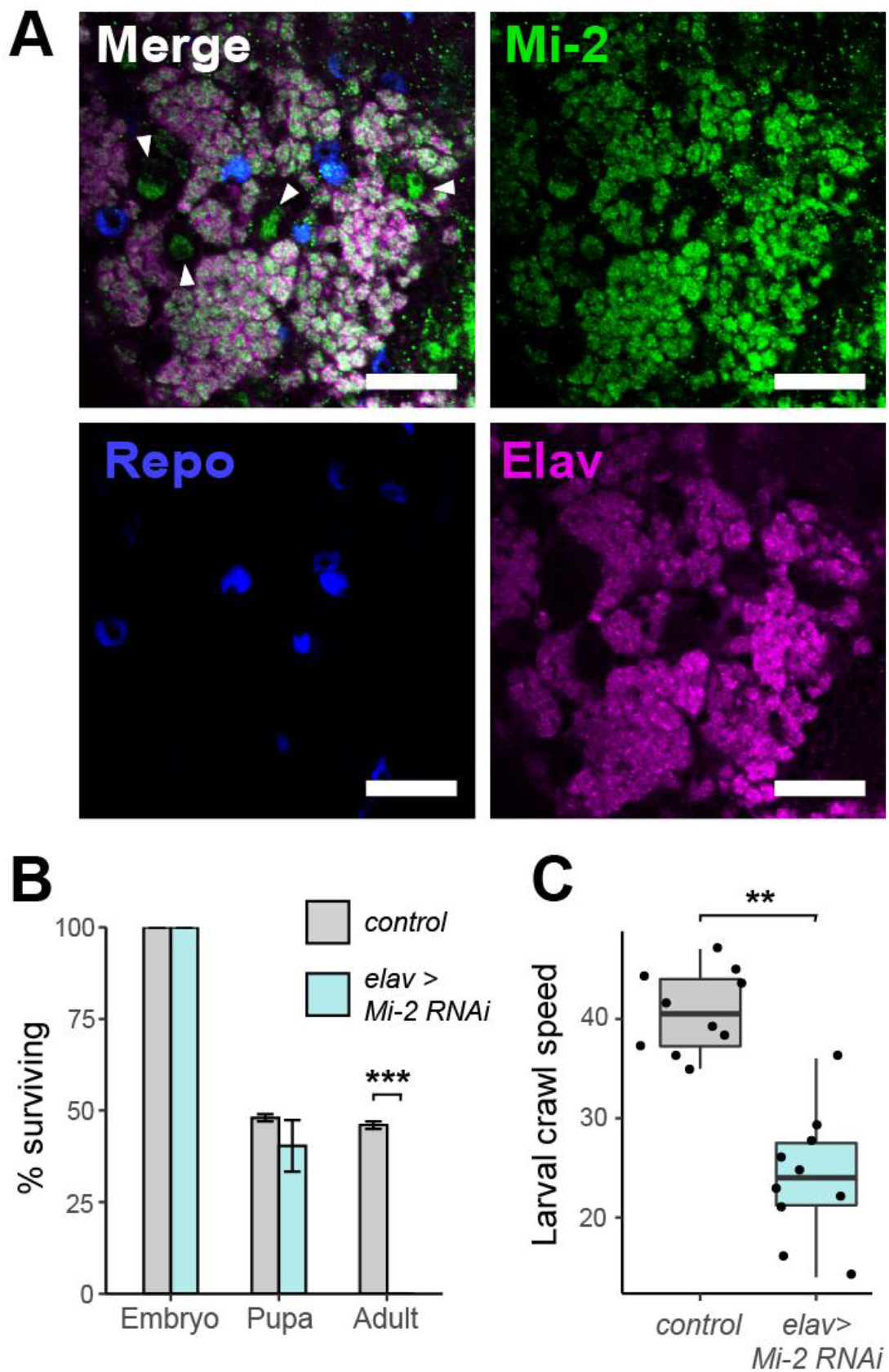
Critical role of *Mi-2* RNAi in *Drosophila* neurons. A) Third instar larval CNS of *Mi-2* GFP-trap line (scale bar = 20 μm). White arrows highlight NSC nuclei. B) Survival of *Mi-2* RNAi and control (*mCherry-RNAi*) flies during development (*** *p < 0.001*). D) Larval crawl speed (cm/minute) in *Mi-2* RNAi and *mCherry* RNAi controls (** *p < 0.01*).

To understand the importance of *Mi-2* expression in neurons, we used RNAi interference (RNAi) to induce cell-type specific knockdown of *Mi-2* in the CNS. An RNAi line targeting Mi-2 was previously shown to produce robust knockdown of Mi-2 in neuronal lineages, therefore, we decided to utilise this line for our study (Zacharioudaki et al., 2019). Using the pan-neuronal *elav-GAL4* driver we observed lethality during late larval/pupal stages (Figure 1B). Whilst neuronal *Mi-2* appeared dispensable for embryogenesis and larval survival (possibly due to maternal contribution of Mi-2), we observed a severe mobility defect in larval crawling compared to controls (Figure 1C) indicating that *Mi-2* activity in neurons is required throughout development at all life-stages for normal neuronal function.

### Mi-2 association with chromatin reveals the presence of distinct complexes

Our data, along with previous reports, suggest that Mi-2/NuRD has an essential role in the regulation of neuronal gene expression. However, it is unclear which genes Mi-2 regulates and whether its function is predominantly via NuRD or dMEC (Mi-2 and MEP-1 only). To better understand NuRD function in regulating neuronal gene expression, we profiled the genomic binding of NuRD complex components using Targeted DamID (TaDa) (Southall et al., 2013). This approach involves fusing a protein of interest to Dam methylase and expressing in a defined cell population, followed by enrichment and sequencing of selectively methylated regions which reflect loci at which the protein associates with DNA (reviewed in (Aughey et al., 2018).

We used Targeted DamID to profile genomic association of Mi-2 as well as NuRD components MTA1-like and HDAC1 in neurons (defined by *elav-GAL4* expression) in the larval CNS. MTA1-like is a core component of the NuRD histone deacetylase subcomplex, with cells lacking MTA orthologues in mammals considered to be effectively NuRD-null (Burgold et al., 2019), whilst HDAC1 is the histone deacetylase subunit (also known as Rpd3). We also assayed for genomic binding of MEP-1, previously reported to associate exclusively with Mi-2 as part of both the dMEC and NuRD complexes. Principal component analysis revealed a single cluster including both Mi-2 and MEP-1 replicates, whilst the remaining NuRD components, MTA1-like and HDAC1, appeared to be distinct (Figure 2A, B and C). Similarity between Mi-2 and MEP-1 binding is further supported by the close correlation between the two datasets (pearson R^2^ = 0.94/0.95), which is almost as strong as that seen between replicates (Figure EV1A). In contrast, both MTA-1like and HDAC1 showed relatively poor correlations to MEP-1 or Mi-2, whereas their correlation with each other was slightly stronger. Whilst HDAC1 binding may be expected to be poorly correlated with Mi-2 due to the inclusion of HDAC1 in protein complexes other than NuRD (e.g. with Polycomb Group proteins (Tie et al., 2001)), the relatively low correlation with MTA1-like points towards the possibility of distinct Mi-2 containing complexes having distinct binding in neurons.

**Figure 2.**
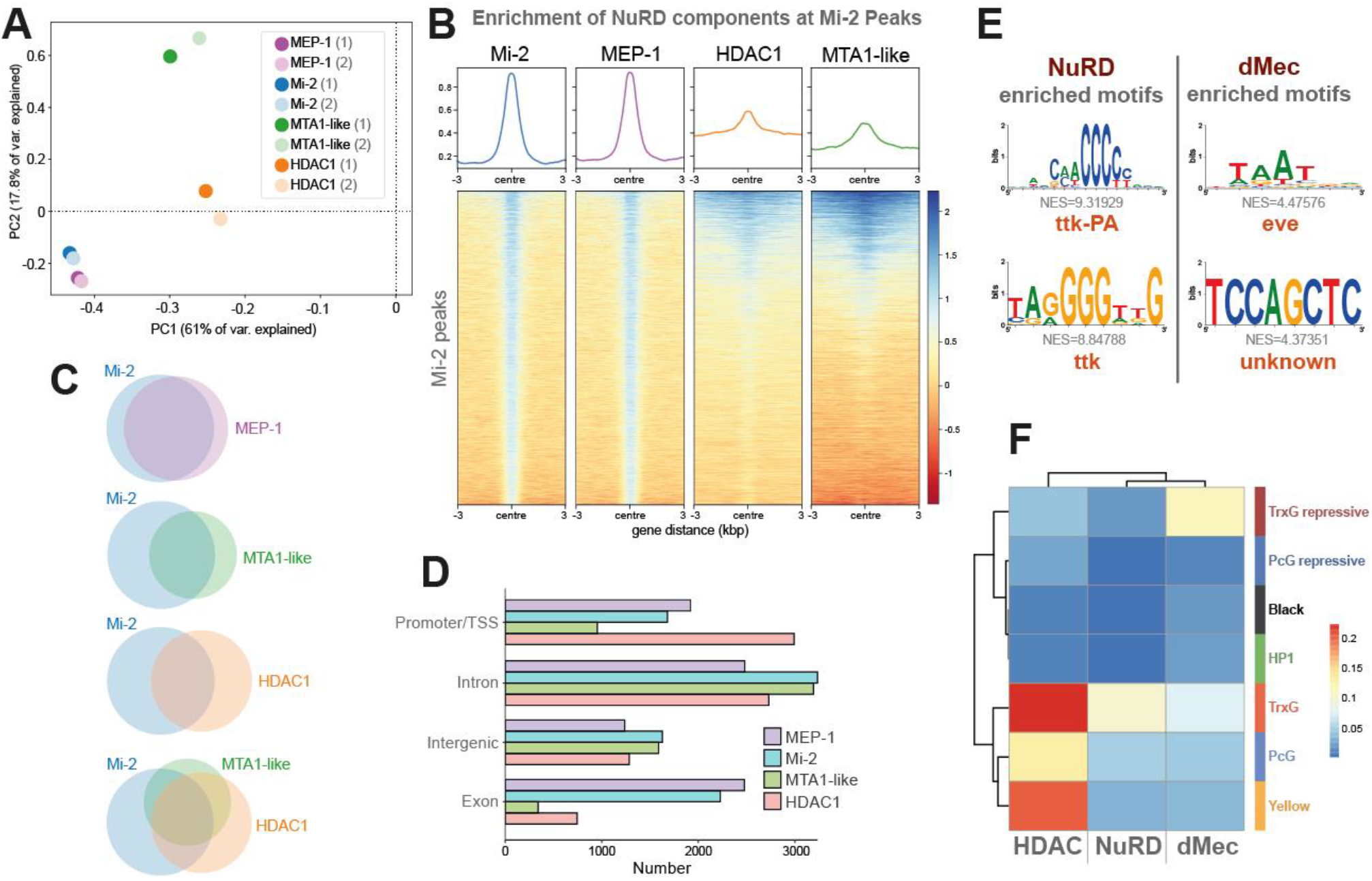
Genomic binding of NuRD components. A) Principal component analysis of all NuRD component binding in neurons including replicates. Mi-2/MEP-1 (blue/purple) cluster independently of MTA1-like (green) and HDAC1 (orange) reflecting greater similarity in their binding profiles. B) Heatmap and enrichment plots indicating binding at Mi-2 peaks. C) Venn diagrams indicating the number of overlapping peaks between NuRD subunits. All intersections between peaks occur at statistically significant frequencies (Fisher’s exact test p < 10^-24^). D) Peak annotation for Mi-2, MEP-1, MTA-1like and HDAC1 peaks. E) Top enriched motifs at NuRD and dMec peaks. F) Heatmap indicating jaccard similarity statistic between chromatin states and NuRD peaks (Mi-2, MEP-1, MTA-1like and HDAC1) and dMEC (Mi-2 and MEP-1 only), or HDAC1 peaks.

To further understand the relationship between Mi-2 and other NuRD components at the chromatin level, Mi-2 significantly bound peaks were compared to those of MEP-1, MTA1-like and HDAC1. We found that the majority of MTA1-like peaks coincided with Mi-2, although a significant number of Mi-2 peaks did not overlap with MTA1-like (Figure 2C). We observed strong binding of HDAC1 throughout the genome, however, a relatively low proportion of HDAC1 peaks intersected with Mi-2 peaks (Figure 2B, C), indicating that Mi-2 is found in association with chromatin without HDAC1 or MTA1-like at the majority of loci.

We observed that around half of Mi-2 peaks coincided with either MTA1-like or HDAC1 with fewer still having all three detected at the same site (around a quarter of total Mi-2 peaks). We also saw that a smaller proportion of HDAC1 peaks were shared with Mi-2 compared to MTA1-like presumably due to the inclusion of HDAC1 in other chromatin modifying complexes (Figure 2C). We also observed some MTA1-like peaks that did not overlap with Mi-2 or HDAC1, suggesting that MTA1-like may have undefined roles outside of NuRD. As previously indicated, MEP-1 and Mi-2 peaks intersected at almost all loci. Inspection of the genomic features at which the various peaks were found to bind demonstrated further disparity between Mi-2/MEP-1 and MTA1-like/HDAC1. We found that whilst HDAC1 was found to be enriched at promoters and TSS, Mi-2 and MEP-1 were comparatively enriched in gene bodies, particularly in exons (Figure 2D). Together these data suggest that multiple Mi-2 containing complexes interact with chromatin in neurons, that may or may not possess histone deacetylase activity. In line with previous reports, we refer to sites bound by Mi-2 and MEP-1 as “dMEC” hereafter (Kunert et al., 2009), whilst sites with significant binding of all four profiled components are considered “NuRD” target sites.

### dMec and NuRD regulate different genes in neuronal differentiation

Examining the gene ontology (GO) enrichment of genes bound by either NuRD or dMec revealed clear differences in the processes that were regulated by each complex. We found that NuRD-regulated genes were enriched for GO terms related to organismal development, with “nervous system development” amongst the top enriched terms (Figure EV1B). In contrast, genes with dMec peaks were enriched for terms relating to cilia function (Lattao et al., 2017) (Figure EV1C) (Appendix Figure S1) reflecting the prevalence of genes that are usually enriched in testis and not expressed in the CNS (although it should be noted that it is possible that some of these genes may also be expressed in fully differentiated sensory neurons). Terms relating to synaptic function were also enriched for genes bound by dMec, which may be expected of a group of genes the expression of which is regulated in neurons. However, inspection of the genes comprising these categories reveals that many of these genes are expressed post-synaptically outside the CNS (*GluRIIA, GluRIIC*) or in later stage pupal CNS in which neurons are in a more differentiated state (e.g. *slo, Syt12*). These results are consistent with a model in which dMEC is largely responsible for repression of gene expression, whilst NuRD is involved in coordinating appropriate transcription of neuronal genes.

We examined NuRD and dMec bound sequences for enrichment of sequence motifs to see if we could identify potential cofactors that may be involved in recruitment or regulation of these complexes. For sequences bound by NuRD, we found that the most highly enriched sequence motifs corresponded well to the consensus sequences for *tramtrack (ttk*) (Figure 2E). ttk has previously been shown to interact with MEP-1 to recruit the NuRD complex (Reddy et al., 2010), and physical interactions with other NuRD components have also been reported (Rhee et al., 2014). In contrast, the *ttk* motif was not enriched for sequences covered by dMec peaks. Instead, the most highly enriched motif identified was a Hox-like motif corresponding most closely to the transcriptional repressor *even-skipped* (*eve*). The differences in the underlying sequences present at NuRD and non-NuRD Mi-2 binding sites suggest that these complexes maybe recruited to target loci by distinct molecular factors.

To gain further insight into how Mi-2 containing complexes regulated gene expression we compared binding to the state of the underlying chromatin. Chromatin can be grouped into discrete states reflecting the occupancy of key regulatory proteins (Filion et al., 2010). We compared the binding of HDAC1, Mi-2, MEP-1 and MTA1-like with published data describing chromatin state domains based on a seven-state model (Marshall and Brand, 2017). We found that HDAC1 binding was associated with broadly permissive chromatin states (Figure 2F). Whilst histone deacetylase activity has been thought to be associated with repression of gene expression, these observations are consistent with previous reports that have described enrichment of HDAC1 binding at active promoters (Wang et al., 2009). Indeed, we also observe strong enrichment of HDAC1 binding at transcription start sites (TSS) as previously reported (Figure EV1D). In contrast, Mi-2, MEP-1 and MTA1-like were found to be associated with a mixture of permissive and repressive states, in particular with Trithorax-like permissive and repressive states (Trx-G / Trx-G_repressive). Taking into consideration only regions that are bound by either Mi-2/MEP-1 (dMec), HDAC1 alone, or all four proteins (i.e. NuRD), we saw that whilst NuRD showed the greatest similarity to the Trithorax-like permissive state, Mi2/dMec more strongly coincided with the transcriptionally silent Trithorax-associated repressive state, which is characterised by the presence of Brahma as well as H1 linker histone. These observations support a model in which NuRD independent Mi-2 binding largely mediates gene silencing/repression, whilst NuRD is involved in the regulation of actively expressed genes.

### *Mi-2* knockdown results in misexpression of non-CNS genes

To determine the effect of Mi-2 loss on gene expression in neurons of the developing CNS, we performed RNA-seq on larval CNS in which *Mi-2* had been knocked down in neurons by RNAi, and compared to controls in which RNAi targeting mCherry was expressed using the same *elav-*GAL4 driver. We found good correlation between replicates (Figure EV2A, B) and observed that the expression of *Mi-2* itself was found to be significantly downregulated (Figure EV2C) in agreement with previous studies (Zacharioudaki et al., 2019). We found that knockdown of *Mi-2* resulted in significantly more upregulated genes than downregulated (397 downregulated vs 1467 upregulated) indicating that in the context of developing neurons, *Mi-2* has a significant role in the repression of gene expression (Figure 3A-C). The top 30 upregulated and top 30 downregulated genes are shown in Figure EV2C, D.

**Figure 3.**
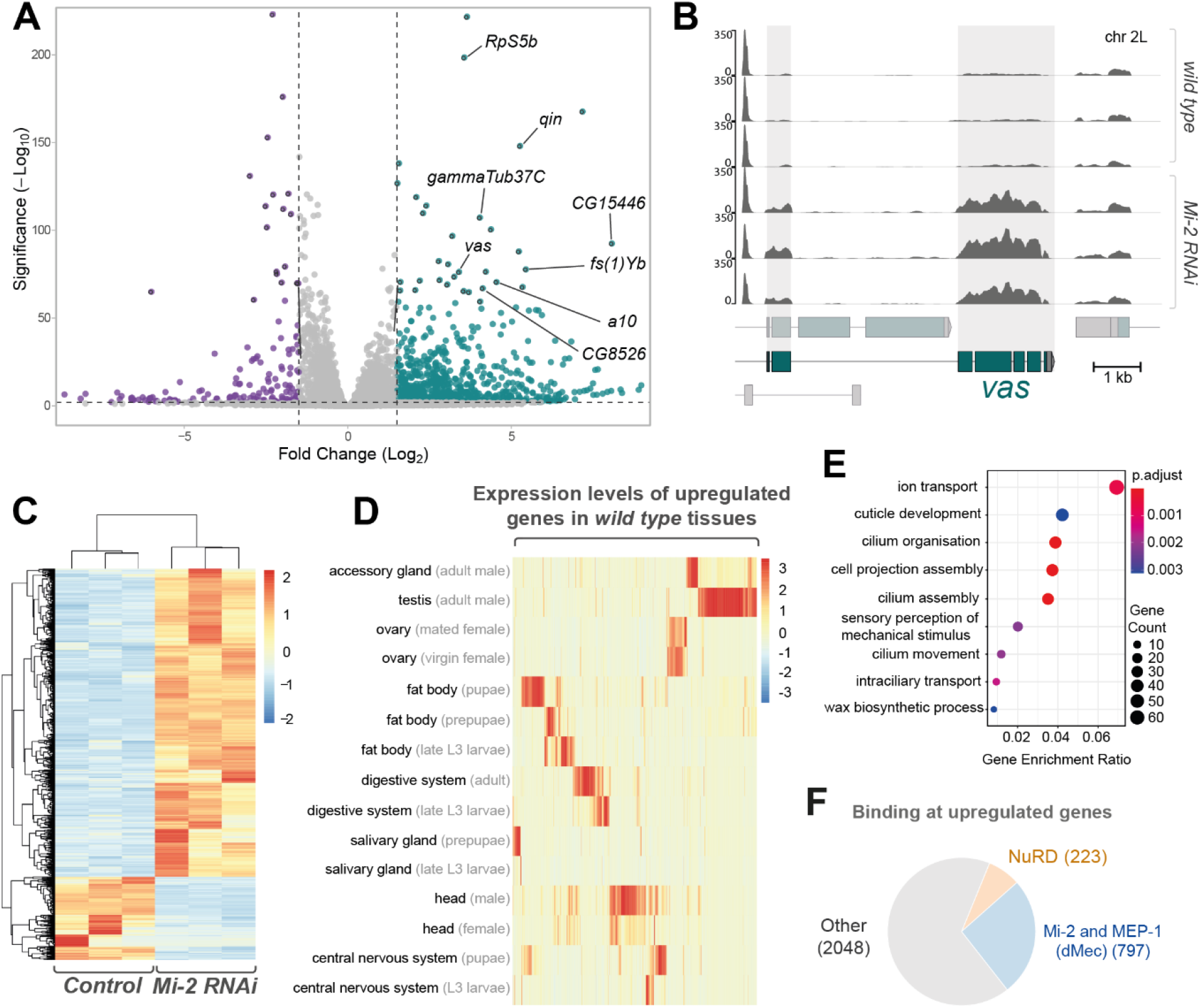
*Mi-2* knockdown in larval neurons results in extensive gene expression changes. A) Volcano plot showing significantly up-regulated (green) and down-regulated (purple) genes with fold-change. Selected mis-expressed genes are annotated. B) RNA-seq tracks showing *vas* mRNA expression in third instar larval CNS (Y-axis is RPKM). C) Heatmap showing relative changes in normalised read-counts between *Mi-2* RNAi and control replicates for all significantly differentially expressed genes (p-adj < 0.05, log2 fold change >1 or <-1). Significantly more genes are upregulated (1467) than downregulated (397). D) Enriched gene ontology terms for genes that are upregulated in *Mi-2* RNAi knockdown. E) Heatmap showing relative expression levels of upregulated genes across selected *Drosophila* tissues (full version in Appendix Figure S2).

Closer inspection of the top up-regulated genes revealed the presence of ectopically expressed genes that are usually expressed in non-CNS tissues such as the germline. For example, *vasa* (*vas*, a well-characterised ovarian germline marker) expression was (~3.5 log2 fold-change) increased compared to controls (Figure 3B). Similarly, *gammaTub37C* – a female germline specific isoform of tubulin, was also very highly upregulated (Figure EV2E), as well as *RpS5b*, encoding a germline specific ribosomal protein (Figure EV2F). We also saw evidence of male-germline specific genes being upregulated, for example *CG15446* (a transcription factor with testis specific expression) was expressed at much higher levels than controls (Figure 3A). Comparison to tissue-specific transcriptomes indicates that the upregulated genes are usually expressed outside the larval nervous system from a broad range of tissues, including a large cluster of genes that have testis-enriched expression, as well as ovary, fat body, and digestive system (Figure 3D and Appendix Figure S2A, B). Furthermore, Gene Ontology (GO) analysis of the significantly up-regulated genes revealed enrichment of GO-terms that may be associated with neuronal maturation or function, such as “ion transport”, “sensory perception” and “cell projection assembly” (Figure 3E). However, we also saw enrichment for unexpected terms such as “cilium assembly” that are not typically associated with nervous system development or function.

We also observed misexpression of neuronal lineage specific genes that are not usually highly expressed in the larval CNS but are expressed in specific neuronal populations in the adult nervous system. For example, the antennal gene *a10* was seen to be very highly upregulated upon knockdown of *Mi-2* in the larval CNS despite usually having expression restricted to the antennae in adult flies (Figure 3A and Figure EV2C). Therefore, these data indicate that Mi-2 is required to ensure fidelity of neuronal transcriptomes by suppressing expression of spatially and temporally inappropriate genes.

### Mi-2/dMEC represses ectopic gene expression independently of HDAC1

Having seen that Mi-2/NuRD components are distributed unequally in the genome, we examined how binding of NuRD components related to differentially upregulated genes. Since previous reports have suggested that dMec has a role in suppressing ectopic gene expression in cultured cells, we questioned whether this complex also plays a greater role in repressing ectopic gene expression in neurons. We found that NuRD binding (i.e. significant binding of Mi-2, MTA1-like, and HDAC1) was found at a much lower proportion of upregulated genes compared to those bound by dMec (Mi-2 and MEP-1) (223 and 797 genes respectively – (Figure 3F)). This observation led us to further investigate the role of dMec in regulating gene expression compared to NuRD.

We used qPCR to confirm upregulation of a selection of highly mis-expressed candidate genes in *Mi-2* knockdown flies (Figure 4A). Since RNAi mediated knockdowns have previously been associated with off-target effects, we repeated these experiments with a second RNAi line which recapitulated the upregulation of ectopic genes previously observed (Appendix figure S3). Since we saw distinct binding of NuRD subunits, we first asked whether *HDAC1* was required for repression of ectopic genes. We found that depletion of *HDAC1* by RNAi in neurons resulted in lethality during pupal stages similarly to *Mi-2* (Appendix figure S4). However, *HDAC1* knockdown with two independent RNAi lines had no effect on expression of any of the genes we tested that were upregulated upon *Mi-2* knockdown. Therefore, whilst neuronal *HDAC1* is essential for survival, HDAC1 is not required for repression of some *Mi-2* induced ectopic gene expression in larval neurons (Figure 4A and Appendix figure S3).

**Figure 4.**
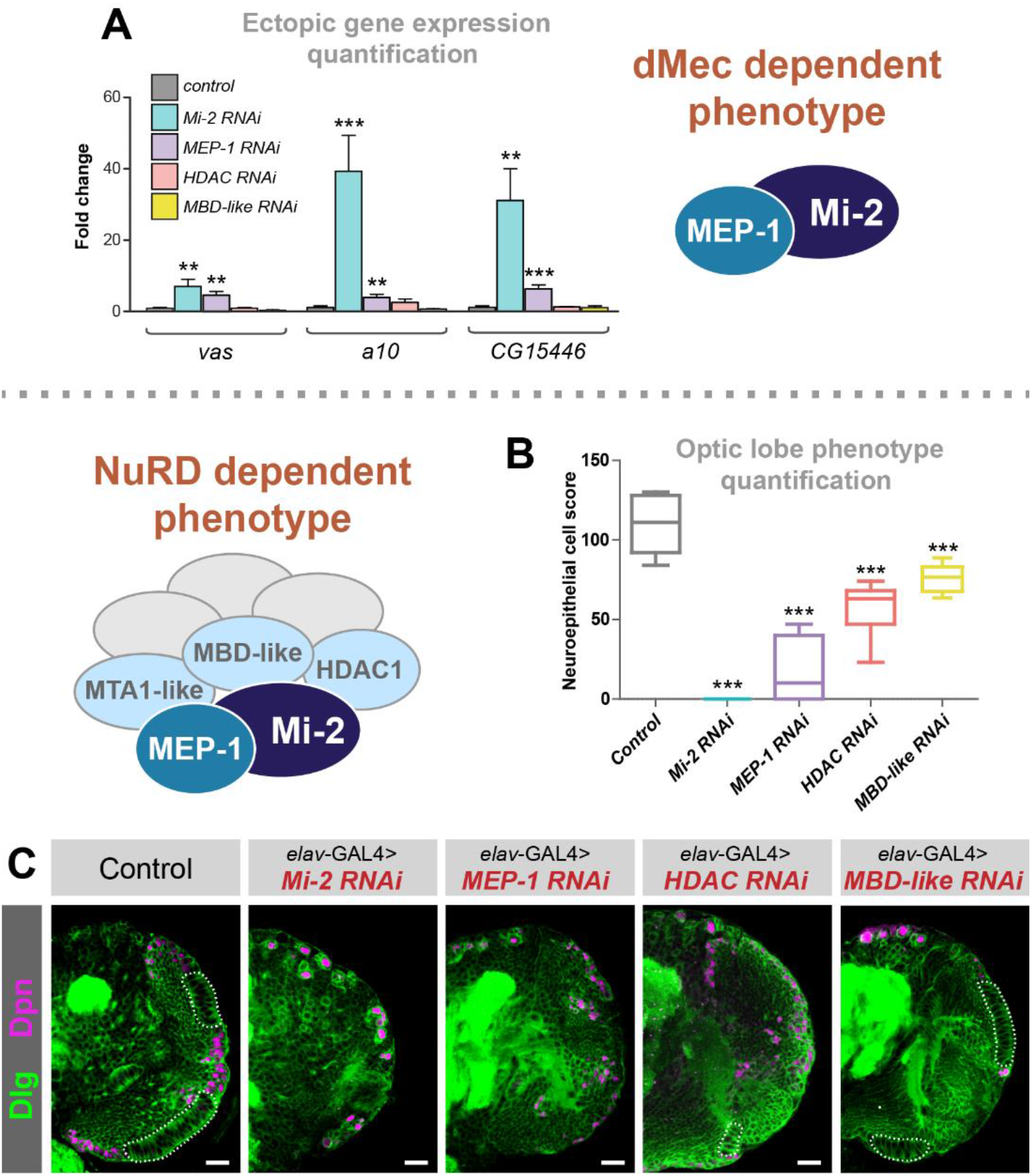
*dMec* knockdown causes ectopic gene expression and *NuRD* knockdown in neurons causes an optic lobe development phenotype. A) Quantification of candidate gene expression (selected from upregulated genes in RNA-seq data) in knockdown genotypes (3 biological replicates for each group). ** *p < 0.01 and ***p < 0.001* (one-tailed student’s t-test or Welch’s t-test against control) Represented as mean ± SEM. B) Quantification of optic lobe neuroepithelial cells of genotypes shown in panel C (n >= 9 for each group). *\*\*\* p < 0.001* (one-tailed student’s t-test or Welch’s t-test against control). Represented as a box plot. C) Representative images of control (*elav-*GAL4*;; mCherry^RNAi^*), *Mi-2* knockdown, *MEP-1* knockdown, *HDAC1* knockdown and *MBD-like* knockdown in the third instar larval CNS (all scale bars = 20 μm).

We questioned whether the expression of ectopic genes in *Mi-2* knockdown was dependent on NuRD (albeit without HDAC1 activity) or mediated by Mi-2 via NuRD-independent mechanisms. To address this, we knocked down *MBD-like*, a core NuRD component thought to bridge the nucleosome remodelling and histone deacetylation subunits. In *MBD-like* RNAi knockdown neurons, no difference in gene expression of ectopic target genes was observed (Figure 4A), providing further evidence of *Mi-2* acting independently of NuRD to mediate gene repression. Since MEP-1 binding in the genome appeared to coincide with Mi-2 at almost every locus, we hypothesised that MEP-1 must also be part of the same complex that confers repression of ectopic gene expression. Consistent with this idea, we saw that knockdown of MEP-1 with *elav-GAL4* was sufficient to cause upregulation of *vas* along with other target genes tested (Figure 4A). Therefore, it appears that dMEC represses the expression of some genes in a MEP-1 and Mi-2 dependent manner, without the requirement for HDAC1-mediated histone deacetylation or other members of the NuRD complex.

### NuRD disruption in larval CNS causes severe optic lobe growth defects

Performing GO analysis for down-regulated genes from our Mi-2 knockdown RNA-seq indicated enrichment of multiple terms related to cell division and stem cell function (Figure EV3A). This is surprising since we knocked down *Mi-2* in neurons, which are exclusively post-mitotic and do not express genes related to cell-division– therefore it is unlikely that Mi-2 could directly downregulate the expression of these genes. To further understand the consequences of *Mi-2* depletion in neurons, we examined dissected third instar larval CNS using confocal microscopy. We found that in *Mi-2* knockdown resulted in severe optic lobe defects. Mi-2 knockdown brains were smaller, and it was evident that all recognisable optic lobe structures (including inner and outer optic proliferation centres as well as differentiating medulla and lamina neurons) were found to be completely absent (Figure 4B, C). Interestingly, this included proliferative cells of the optic lobes and neuroepithelium (NE), including the neuroblasts. We also examined *Mi-2* knockdown ventral nerve cords (VNCs) and observed no morphological changes or differences in NSC numbers (Figure EV3B, C). Therefore, we attributed the enrichment of GO terms relating to precursor cell function to the fact that a large number of cells expressing these genes were not present in the tissue from which RNA was extracted. Due to this severe phenotype, in which a large fraction of optic lobe cells are absent, we decided that we could not reliably identify genes that were downregulated as a consequence of Mi-2 loss rather than due to absence of cells expressing those genes. However, since we still saw far fewer downregulated than upregulated genes, this reinforces our conclusion that Mi-2 mainly acts to suppress gene expression in this context.

Having seen NuRD-independent effects of *Mi-2* on gene expression, we questioned whether knockdown of other NuRD complex components would phenocopy the optic lobe phenotype we observed when *Mi-2 was* knocked down with *elav-GAL4*. In *MEP-1, HDAC1* and *MBD-like* knockdowns, the larval CNS displayed a similar optic lobe defect to that of *Mi-2* (Figure 4B,C). Quantification of NE cells indicated *that MEP-1, HDAC1* and *MBD-like* knockdowns had significantly fewer NE cells (Figure 4C), however the phenotype was less severe with a small number being present in most brains (although this may be due to less efficient RNAi mediated knockdowns in these lines). (Burgold et al., 2019) We also observed an optic lobe phenotype with knockdown of *MTA-1* (Figure EV3D, E). Therefore, due to the similarity of phenotypes observed with knockdowns of multiple NuRD complex members we concluded that whilst *Mi-2* functions to repress some ectopic genes in a NuRD independent manner, its effect on optic lobe development involves the activity of the NuRD holocomplex (although it remains unclear whether dMEC also contributes to this phenotype). Conversely, the ectopic upregulation of genes is independent of NuRD/histone deacetylation, and is not required to produce optic lobe developmental defects in the larva (although since the phenotypes vary in severity, there may be some contribution of dMec induced ectopic gene expression on NE survival).

### Fully differentiated neurons do not require Mi-2 for silencing of ectopic gene expression or proper optic lobe development in larvae

Factors involved in the maintenance of neuronal transcriptomes during early neuronal maturation can have less severe phenotypes in fully differentiated neurons in which synapses have been formed (Hassan et al., 2020; Southall et al., 2014). This correlates with observed changes to the underlying chromatin landscape which is dynamic during differentiation and neuronal maturation (Aughey et al., 2018; Marshall and Brand, 2017). Notch-induced hyperplasia in neuronal lineages is enhanced by the knockdown of *Mi-2*, however, this effect is diminished as cells become more differentiated (Zacharioudaki et al., 2019). In our previous knockdown experiments, we employed the *elav-*GAL4 driver to target all neurons in the *Drosophila* larval CNS. This includes a substantial proportion of neurons that are post-mitotic cells, but not fully differentiated to the point at which synaptic proteins are expressed. In fact, it has been suggested that the majority of larval brain *elav*-expressing neurons contribute relatively little to neuronal function (Eichler et al., 2017), since they do not possess axons or dendrites or express ion-channels associated with neurotransmission (Ravenscroft et al., 2020), despite comprising a large proportion of the total cells in the larval brain (Jiao et al., 2021). Therefore, we questioned whether Mi-2 dependent optic lobe phenotypes or de-repression of silenced genes would be observed when Mi-2 was knocked down in either progenitor cells or fully differentiated neurons.

The larval CNS consists of mostly immature neurons that are produced post-embryonically. However, there is a sizeable population of fully differentiated neurons that persist from the embryo that express the mature neuron marker nSyb. The majority of these are found outside of the optic lobes in the central brain and VNC, however there are a very small number of nSyb cells found in the optic lobe lamina (Jiao et al., 2021). To rule out the contribution of signalling from these mature neurons being responsible for the morphological phenotypes, we examined brains in which *Mi-2* RNAi was driven with *nSyb-GAL4*. Knockdown of Mi-2 using nSyb-GAL4 did not result in any detectable change in *vas* mRNA levels (Figure 5A). We also tested whether depletion of Mi-2 in adult neurons was sufficient to de-repress *vas* expression using a temperature sensitive GAL80 to de-repress *Mi-2* RNAi in neurons using *elav*-GAL4 for a 24-hour period after eclosing. As in third instar larvae, no change in *vas* expression was detectable in adult heads (Figure EV4A). We also tested for expression of *a10* and *CG15446*, which also did not show any increased expression in adult neurons, indicating that Mi-2 is only required for the repression of ectopic gene expression in the early stages of neuronal development. We used *wor-*GAL4, to drive *Mi-2* RNAi in NSCs (it should be noted that this driver is likely to also affect the newly born GMC and early differentiating neurons due to perdurance (Johnson et al., 2018) (Figure 5B). Using *wor-*GAL4 to knock down *Mi-2* in NSCs, we performed qPCR to detect changes in gene expression. When *Mi-2* was knocked down in NSCs, the expression of female germline marker *vas* was significantly upregulated (Figure 5A), as previously seen with *elav-*GAL4 knockdowns. *vas* expression was upregulated to a lower extent than when induced with *elav-*GAL4 (~4 fold increase compared to ~10 fold), however this may be due to the lower number of affected cells expressing *wor-*GAL4 rather than reflecting a difference in the requirement for Mi-2 for repression of *vas* in NSCs. It should also be noted that it is possible that a small amount of perdurance of GAL4 into NSC progeny may be responsible for de-repression of non-neuronal target genes in immature neurons or GMCs, rather than due to an effect exclusively in NSCs.

**Figure 5.**
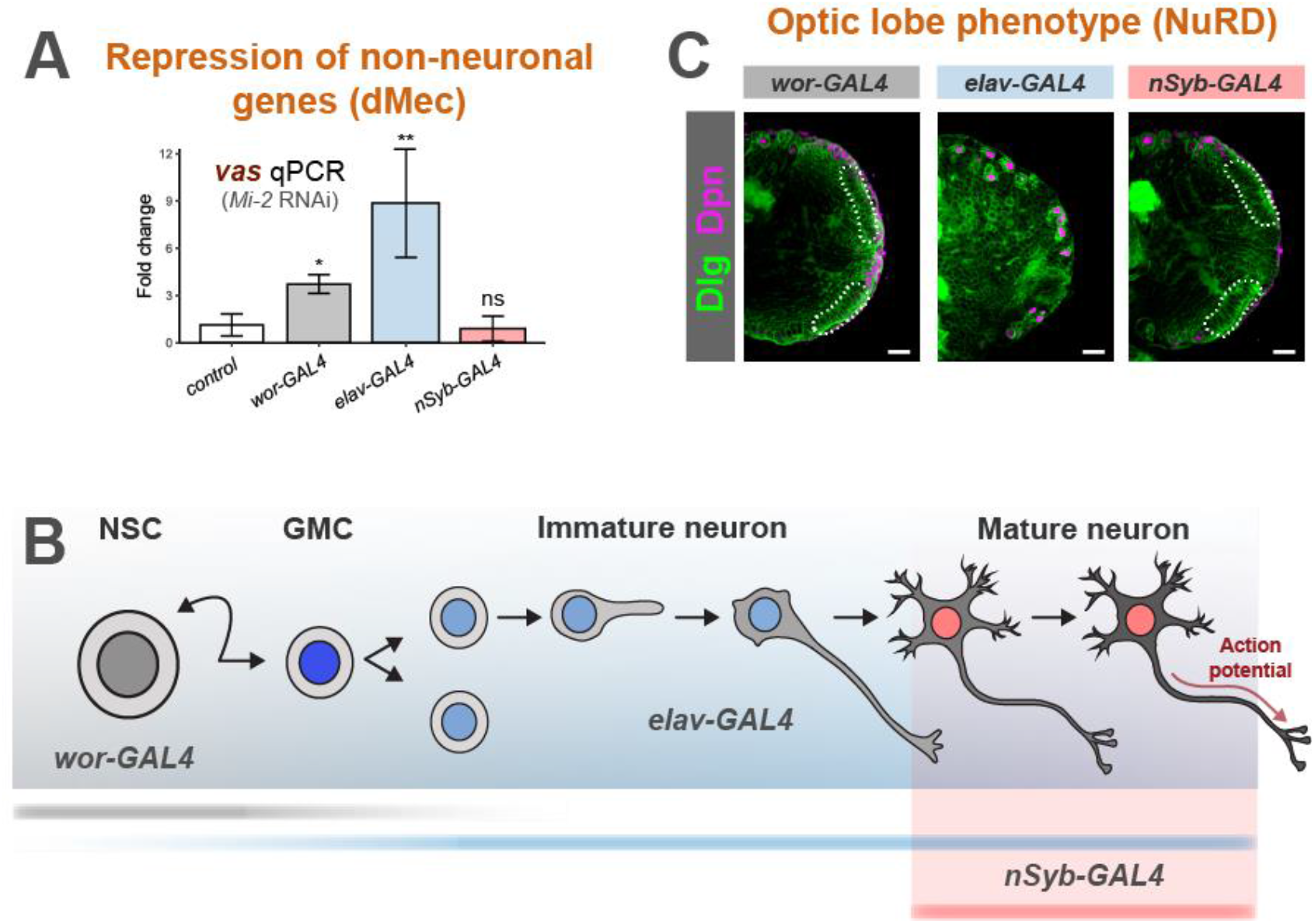
Differential requirements of Mi-2 during neuronal differentiation. A) Ectopic expression of *vas* when *Mi-2* is knocked down at different stages of differentiation. (3 biological replicates for each group). *\*p < 0.05* and ** *p < 0.01* (one-tailed student’s t-test against control) Represented as mean ± SEM. B) Representation of expression of GAL4 drivers used. C) Larval brain optic lobe morphology when *Mi-2* is knocked down at different stages of differentiation Elav = green, Dpn = magenta.

Surprisingly, we found that *Mi-2* knockdown in NSCs showed no apparent defects in optic lobe development, with brain lobes having a similar morphology and neuroepithelial cell numbers to controls (Figure 5C and Figure EV4B, C). In contrast to when *Mi-2* was depleted in neurons (Figure 1D), loss of *Mi-2* in NSCs did not result in larval locomotion defects (Figure EV4D). However, *Mi-2* knockdown in NSCs did result in lethality, with 100% dying as pupae. Therefore, since brain size, morphology and movement are normal in these animals, these results suggest that Mi-2 is not required for stem cell division or differentiation into neurons. Nevertheless, Mi-2 function in NSCs is required for the correct functioning of subsequently born neurons.

Having seen that expression of *Mi-2* RNAi in neuronal precursors was not responsible for the optic lobe phenotype, we questioned whether other unexpected expression may be responsible. *elav-*GAL4 may drive some expression in embryonic glial cells (Berger et al., 2007), therefore, we questioned whether unexpected glial knockdown of *Mi-2* may have resulted in a disruption of signalling that maintains the optic lobe stem cells. To address this, we repeated the *Mi-2* knockdown with an *elav-*GAL4 driver combined with *repo-GAL80*, to repress unwanted expression of the RNAi hairpin in glia. In these animals, we observed the same severe optic lobe phenotype as previously with *elav-*GAL4 alone (Figure EV4E). These data indicate that *Mi-2* knockdown in neurons may affect optic lobe growth via a non-cell-autonomous mechanism. This may be partly explained by the requirement for innervation by photoreceptor neurons to initiate neurogenesis in the developing visual centres (Huang and Kunes, 1996; Selleck and Steller, 1991), which would be affected by *elav-*GAL4 knockdown of *Mi-2*. We also examined 1^st^ instar *Mi-2* knockdown brains and found that the typical rosette of NE cells is present (Figure EV4F). Therefore, the NE is present at earlier stages of development, however, is not persisting to later stages. As *elav*-GAL4 has never been reported to be expressed in NEs, the role of any glial expression of *Mi-2* RNAi has been eliminated (Figure EV4E), and knockdown of *Mi-2* in neuroblasts has no phenotype (Figure EV4B, C), this strongly suggests that there is a non-cell-autonomous mechanism occurring. However, a full explanation of this phenotype is beyond the scope of this study and further work must be conducted to determine the mechanism by which the optic lobe defect occurs.

In contrast to knockdown in all neurons including immature neurons with *elav-*GAL4, depletion of *Mi-2* exclusively in mature neurons did not produce overt morphological phenotypes and no significant difference in neuroepithelial cell numbers were detected when compared to controls (Figure 5C and Figure EV5A, B). As with the other drivers used, *Mi-2* knockdown with *nSyb-GAL4* resulted in 100 % lethality during pupal stages. Therefore, whilst Mi-2 is required in mature neurons, the morphological defects observed in the larvae do not arise from loss of Mi-2 function in fully differentiated neurons in the larval CNS.

### Dynamic Mi-2 binding during neuronal differentiation

We reasoned that varying consequences of *Mi-2* loss at different stages of neuronal differentiation may be reflected by changes to Mi-2 binding in different cell types. To examine differences in Mi-2 genome occupancy in precursor cells vs fully differentiated cells, we profiled Mi-2 (using TaDa) in two further populations of cells in the larval brain. Firstly, in NSCs (*wor*-GAL4 - This driver has been used previously to produce Targeted DamID profiles that are distinguishable from those produced from GMCs/newly born neurons. However, it should be noted that due to perdurance of GAL4, these profiles likely represent the early stages of neurogenesis more generally rather than specifically NSCs. We then also profiled mature neurons (*nSyb*-GAL4) in the larval CNS, in which we saw no change in *vas* expression upon *Mi-2* knockdown. Since these neurons do not reflect the direct descendants of those produced by the *wor*-expressing NSCs in the larval brain (being mostly derived from earlier embryonic neurogenesis), they may also display lineage specific changes in Mi-2 binding.

Binding profiles between cell types were compared and regions of differential binding identified (Figure 6 A,B). Total neurons (*elav-*GAL4) show the largest number of significantly bound peak regions (10,420), with mature neurons binding 7054 genes (Figure 6B). However, the number of peaks common to each cell type is relatively low. Therefore, Mi-2 binding is highly dynamic during neuronal differentiation. In many individual genes, sites with enriched binding in NSCs showed a different set of enriched Mi-2 binding sites in immature neurons (Figure 6 A, C). A similar change occurs from immature to mature neurons, with 2144 genes that contain both immature-vs-mature and mature-vs-immature regions of Mi-2 enrichment (Figure 6C and the example of *chn* in Figure 6A).

**Figure 6.**
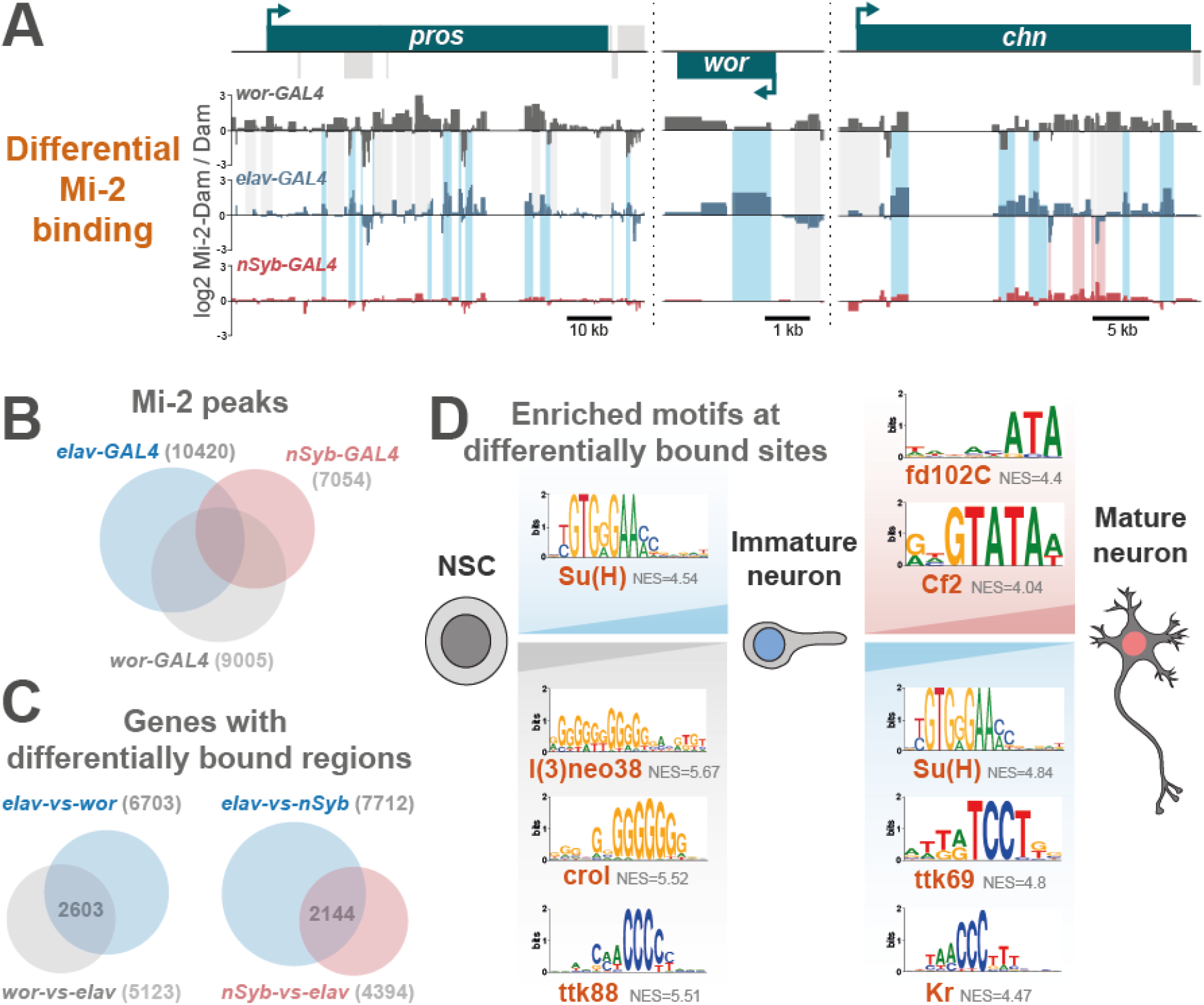
Dynamic Mi-2 binding during neuronal differentiation. A) Examples of differential genomic binding of Mi-2 at neural genes. B) Overlap of Mi-2 binding peaks in NSCs, immature neurons and mature neurons. C) Number and overlap of genes with differential Mi-2 binding. D) Motif analysis of differentially bound regions.

Motif enrichment analysis of the differentially bound regions identified transcription factor binding sites (Figure 6D). Binding sites for l(3)neo38, crol and ttk88 are present in the NSC-vs-immature comparison, suggesting that these sites are occluded in NSCs but available for binding in neural progeny. Interestingly, l(3)neo38 occupies a set of putative neuronal enhancers in the developing embryo and is required for the proper formation of the larval brain (Reddington et al., 2020). l(3)neo38 and crol have both been implicated in the regulation of heterochromatin silencing (Schneiderman, 2010 & Swenson, 2016). While ttk88 sites are enriched for binding in NSC-vs-immature neurons, ttk69 sites (a different splice isoform) are enriched in the immature-vs-mature comparison. Finally, fd102C and Cf2 sites have greater Mi-2 binding in mature-vs-immature neurons. Fd102C is an uncharacterised forkhead transcription factor highly expressed in the developing and adult nervous system (Krause et al., 2022) and Cf2 is a zinc-finger factor involved in follicle cell fate and myogenic gene expression (Bagni et al., 2002; Hsu et al., 1996). These data indicate that a dynamic repertoire of transcription factors may direct or be influenced by dynamic Mi-2/NuRD binding during neuronal maturation. It is clear from the broad changes in binding that Mi-2 coordinates a distinct gene expression programme in early neuronal lineages compared to fully differentiated neurons corresponding to the differences in phenotypic outcomes observed (although some of the observed changes may be due to embryonic vs larval neuronal lineage differences).

Su(H) (Notch co-factor) motifs were enriched in both immature-vs-NSC and immature-vs-mature regions (Figure 6D). This aligns with previous work showing that Mi-2 is required to deactivate notch responsive stem-cell enhancers in neuronal progeny (Zacharioudaki et al., 2019). Mi-2 was implicated in the silencing of genes in the *E(spl)-C* locus, with broad binding of Mi-2 across this locus having been demonstrated in *Drosophila* cell lines (Ho et al., 2014; Kreher et al., 2017). Consistent with this result we saw strong binding of Mi-2 across the *E(spl)-C* locus in both NSCs and immature neurons (Figure EV5C). This study also showed that in more mature neurons Mi-2 depletion was not sufficient to enhance notch-induced overgrowth. We saw no significant binding of Mi-2 in mature neurons across the entire region (approx. 50 kb), supporting the conjecture that in more fully differentiated cells, continued chromatin remodelling of the *E(spl*) genes by Mi-2 is not required for their repression.

## Discussion

Our data provide evidence for Mi-2 activity in repressing ectopic gene expression independently of NuRD. Mi-2 from *Drosophila* or humans is sufficient for the remodelling of nucleosomes *in vitro*, in the absence of other NuRD components (Brehm et al., 2000; Wang and Zhang, 2001). In mouse embryos, Mi-2 orthologue CHD4 was shown to be necessary for lineage specification in a manner that was independent of MBD proteins that are required for a functional NuRD complex (O’Shaughnessy-Kirwan et al., 2015). Similarly, loss of CHD4 in human erythroid cells resulted in de-repression of fetal *γ-globin*, which occurred independently of NuRD (Amaya et al., 2013). Furthermore, similarly to what we observe in neurons, MBD3 was found to bind at less than half of all CHD4 sites in mouse ESCs and were anti-correlated to histone markers associated with transcription (Bornelov et al., 2018). Therefore, it is likely that NuRD-independent Mi-2 regulation of gene expression is a conserved feature in animal cells.

A previous study concluded that Mi-2 in complex with MEP-1 (dMec) was the dominant Mi-2 containing complex in *Drosophila* Kc cells (Kunert et al., 2009). From this observation it could be inferred that NuRD plays a relatively minor role in the regulation of gene expression. However, this conclusion was drawn based on biochemical interactions and relative stoichiometries of NuRD components. We find that in contrast to the abundance of dMec, around a third of Mi-2 peaks intersect with MTA1-like, indicating that NuRD is prevalent at a large number of loci. Furthermore, severe optic lobe phenotypes were observed with multiple NuRD components, which did not always include de-repression of lineage-inappropriate genes. These data indicate that despite being present at lower amounts than dMec, NuRD plays a critical role in nervous system development. Whilst it is possible that the proportions of NuRD/dMec may differ between the CNS and cultured cells, our data suggest that the relative amounts of protein detected by biochemical means may not be reflective of either the distribution of sites at which complexes are associated in the genome, or the severity of consequences resulting from their loss. It is currently unclear whether an analogous complex to dMEC exists in mammals. MEP-1 does not share direct orthology with any single mammalian transcription factors, however, since it is clear that CHD4 is capable of conferring NuRD-independent activities in mammalian cells, it is possible that there are unidentified mammalian genes that act in an analogous role to MEP-1.

Our data indicates that Mi-2 represses non-neuronal genes such as those normally expressed in the germline, and that this is largely independent of NuRD. Interestingly, Mi-2/dMec depletion resulted in proneural gene expression in Kc cells (Kunert et al., 2009), suggesting that Mi-2 may repress ectopic gene expression in multiple tissue types dependent on context. A similar phenotype has also been reported in *C. elegans* in which MEP-1/Mi-2 knockdown resulted in expression of germline genes in somatic tissues, indicating that this may be a conserved function of Mi-2 (Unhavaithaya et al., 2002). It is possible that in mammals, NuRD-independent CHD4 function is also required for gene repression since loss of CHD4 in lymphocytes results in upregulation of non-hematopoietic genes (Arends et al., 2019). However, it remains unclear whether this transcriptional repression is dependent on histone deacetylase activity/NuRD, or NuRD-independent CHD4 activity.

Mi-2 is only required to suppress ectopic gene expression up to the early stages of neuronal development, whereas in more fully differentiated cells it is dispensable. Therefore, it is likely that other molecular factors are required for the repression of these genes following synaptogenesis. Mi-2/dMEC may be required to establish a stable epigenetic state following recruitment by sequence specific transcription factor binding, after which these states are maintained by separate chromatin modifying proteins or complexes. The degree of differential binding of Mi-2 observed during neuronal differentiation is consistent with the idea that Mi-2 coordinates a distinct gene expression programme during early neuronal differentiation compared to in mature neurons. The identification of different transcription factor binding sites during development suggests that either the cognate transcription factor is recruiting Mi-2 to these sites, or that Mi-2 is independently positioned there to establish a chromatin state to prevent the binding of the factor. Since it seems that Mi-2 exhibits at least two independent modes of interacting with chromatin (i.e. via dMEC/NuRD), either or both of these models may be true depending on the presence of other subunits and wider genomic context. Su(H) binding sites are found at sites enriched in immature neurons versus both NSCs and mature neurons (Figure 6D). Furthermore, l(3)neo38 binding sites, that have been identified at neural enhancers (Reddington et al., 2020), are more strongly bound in NSCs compared to immature neurons. Therefore, for at least Su(H) and l(3)neo38, Mi-2 appears to be blocking binding of these factors (loci specific) at stages of differentiation where their function is not required/wanted.

Together, our data highlight that the activity of Mi-2 is dependent on both its molecular and developmental context. These data highlight the importance of detailed analyses of individual subunits in inferring functions of chromatin modifying complexes in complex multicellular tissues to fully understand their role in gene regulation.

## Materials and Methods

### Fly stocks

Unless stated otherwise flies were kept on standard fly food media at 25 degrees. A full list of fly stocks used in this study can be found in Appendix Table S1. Mixed sex larvae/flies were used for all experiments.

### Immunohistochemistry and imaging

Third instar larval CNS were dissected in PBS and fixed for 20 min with 4% formaldehyde in PBS. The following antibodies were used in this study: rat anti-Elav (7E8A10 - Developmental Studies Hybridoma Bank), chicken anti-GFP (600-901-215 - Thermo scientific), mouse anti-DLG (4F3 - Developmental Studies Hybridoma Bank), and guinea pig anti-Dpn (kind gift from A. Brand). Samples were imaged using a Zeiss LSM510 confocal microscope and images processed using LSM Image Browser and image J. To quantify an OPC NE cell score for each genotype, 3^rd^ instar larval brains were mounted dorsal side down and full image stacks (2 μm intervals) of the brain lobe were taken using the DLG channel. The number of OPC NE cells were counted at image slices at ¼, ½ and ¾ through the stack (identified by position and morphology) and added together to give the score. To quantify the number of Dpn positive cells in VNCs,, whole VNC image stacks were analysed using the 3D Objects Counter in Fiji (Bolte and Cordelieres, 2006). For each genotype, a Shapiro–Wilks test was performed to calculate if the experimental and control data had normal distributions. If they did, a one-tailed student’s t-test was applied, and if not a Welch’s t-test was applied.

### Generation of Targeted DamID lines

*MEP-1* and *Mi-2* were amplified by PCR from cDNA with primers including 30 bp complementary sequence to *pUASTattB-LT3-Dam* (Southall et al., 2013) (see Appendix Table S2). Linearised *pUAST-LT3-Dam* (digested with NotI and XbaI) was combined with either *MEP-1* or *Mi-2* PCR-amplified coding sequences in a Gibson assembly reaction using NEBuilder^®^ HiFi DNA Assembly kit (NEB) following manufacturers protocols. Resulting N-terminal fusion constructs were injected into *attP2* flies (*y w P{y[+t7.7]=nos-phiC31\int.NLS}X #12;; P{y[+t7.7]=CaryP}attP2* – Department of Genetics Microinjection Service, Cambridge) for integration on the third chromosome.

*UAS-MTA-1-like-Dam* and *UAS-HDAC1-Dam* were generated using the FlyORF-TaDa cassette exchange protocol as previously described (Aughey et al., 2021). Briefly, *UAS-HDAC1* and *UAS-MTA1-like* FlyORF lines were crossed to *hsFLP; TaDa-FlyORF*. Progeny were heat-shocked at 37 degrees for 45 mins at 96 hours following larval hatching. Resulting males were crossed to *Dr/TM6B* and white-eyed F2 males and female virgins used to create a stable stock.

### Targeted DamID

Dam fusion or Dam-only males were crossed to virgins of the appropriate driver line for expression of Dam in the desired cell-types. For expression in NBs, all larval neurons (including immature neurons), or mature neurons only - the drivers *wor-*GAL4*; tub-GAL80^ts^, elav-GAL4; tub-*GAL80^ts^, *and nSyb-GAL4; tub-*GAL80^ts^, were used, respectively. Embryos were collected for 4 hr then raised at 18°C to prevent premature Dam expression. Animals were transferred to 29°C at 8 days after embryo deposition for 24 hours. 60 larval central nervous systems were dissected and stored at −80 degrees. Genomic DNA extraction, methylated DNA enrichment and library preparation were carried out as previously described (Marshall et al., 2016). Two biological replicates were generated for each experiment.

Libraries were sequenced with 50 bp single end sequencing on Illumina HiSeq or NovaSeq platforms. >15 million reads were obtained for each library. Data were processed using a previously described Perl pipeline (Marshall and Brand, 2015) (https://github.com/owenjm/damidseq_pipeline).

Peak calling was performed using a previously described Perl program (available at https://github.com/tonysouthall/Peak_calling_DamID) which allows for the identification of broadly bound regions that characterise DamID data (Estacio-Gomez et al., 2020). Briefly, false discovery rate (FDR) was calculated for regions consisting of >1 GATC bounded fragments for each replicate. Significant (FDR<0.01%) regions present in all replicates are merged to form a final peak file. Enriched motifs discovery was performed using i-cisTarget on significant peak regions (Herrmann et al., 2012). Intersecting regions for comparison of binding between Mi-2, MEP-1, MTA1-like, or HDAC1 were computed using bedtools intersect utility (Quinlan and Hall, 2010). Similarity between binding peaks was determined using the bedtools jaccard utility, which calculates the ratio between the number of intersecting base pairs and the union between two coordinate sets (Favorov et al., 2012).

### Larval crawling assays

Larval crawling assays were conducted as in (Nichols et al., 2012). Briefly, individual third instar wandering larva were placed on room temperature agar with a 10 mm^2^ grid placed underneath as a distance reference. Larvae were left to acclimatise for 1 min after which larval movement was tracked visually. Ten animals were used per genotype with three technical replicates performed per animal.

### Developmental survival assays

Flies of the appropriate genotype were left in cages containing standard food media supplemented with yeast and left to acclimatise over three nights at 25 °C. Food media was exchanged for apple juice agar and flies left to lay for 5 h. Following the laying period, 100 embryos were transferred to fresh food vials to allow for counting of surviving adults and pupae.

### RNA extraction

Three biological replicates were used for knockdown or control samples. Third instar larval CNS were dissected from thirty animals per replicate and total RNA extracted using TRIzol reagent (https://tools.thermofisher.com/content/sfs/manuals/trizol_reagentpdf) following suppliers protocols. Library preparation and sequencing were subsequently performed by Beijing Genomics Institute (BGI). Sequencing data was mapped to the *Drosophila* genome (release 6.22) using STAR (Dobin et al., 2013). Mapped files were collected in a matrix using featureCounts from the Rsubread package (Liao et al., 2019). Differential expression analysis and MA plots were carried out using the Deseq2 R package (Love et al., 2014).. Genes that had an adjusted p-value <0.05 and a log2 fold change greater than 1 (for upregulated) or less than −1 (for downregulated) were classified as significant. Heatmap generated using pheatmap package in R. Analysis of tissue specific gene enrichment was performed using DGET (Hu et al., 2017).

### qRT-PCR

Following RNA extraction, reverse transcription was performed using the iScript cDNA synthesis kit (Bio-Rad), following manufacturer’s protocols. qRT-PCR was carried out using iTaq Universal SYBR Green Supermix (Bio-Rad). Expression of target genes was normalised to *Ribosomal Protein L4* (*Rpl4*) and compared to expression in control samples expressing RNAi targeting *mCherry*. Primers amplifying sequences <200 bp were designed to span exon junctions where possible (sequences used can be found in Appendix Table S2. Three biological replicates were carried out for each genotype, with at least two technical replicates for each biological replicate. For each genotype, a Shapiro–Wilks test was performed to calculate if the experimental and control data had normal distributions. If they did, a one-tailed student’s t-test was applied, and if not a Welch’s t-test was applied.

## Supporting information

Expanded view figures

## Data availability

All raw sequence files and processed files have been deposited in the National Center for Biotechnology Information Gene Expression Omnibus (GSE199146).

## Acknowledgments

We would like to thank the Southall lab and Imperial College fly community for help and feedback on this project. We would like to thank Andrea Brand for generously providing fly stocks and the Dpn antibody. We also thank the Bloomington *Drosophila* Stock Center (NIH P40OD018537) for fly stocks. This work was funded by a Wellcome Trust Investigator grant 104567 to T.D.S and a BBSRC grant BB/P017924/1 to T.D.S. and G.N.A.

## References

Amaya, M., Desai, M., Gnanapragasam, M.N., Wang, S.Z., Zu Zhu, S., Williams, D.C., Jr., and Ginder, G.D. (2013). Mi2beta-mediated silencing of the fetal gamma-globin gene in adult erythroid cells. Blood 121, 3493–3501.

Arends, T., Dege, C., Bortnick, A., Danhorn, T., Knapp, J.R., Jia, H., Harmacek, L., Fleenor, C.J., Straign, D., Walton, K., et al. (2019). CHD4 is essential for transcriptional repression and lineage progression in B lymphopoiesis. Proc Natl Acad Sci U S A 116, 10927–10936.

Aughey, G.N., Delandre, C., McMullen, J.P.D., Southall, T.D., and Marshall, O.J. (2021). FlyORF-TaDa allows rapid generation of new lines for in vivo cell-type-specific profiling of protein-DNA interactions in Drosophila melanogaster. G3 (Bethesda) 11.

Aughey, G.N., Estacio Gomez, A., Thomson, J., Yin, H., and Southall, T.D. (2018). CATaDa reveals global remodelling of chromatin accessibility during stem cell differentiation in vivo. Elife 7.

Bagni, C., Bray, S., Gogos, J.A., Kafatos, F.C., and Hsu, T. (2002). The Drosophila zinc finger transcription factor CF2 is a myogenic marker downstream of MEF2 during muscle development. Mech Dev 117, 265–268.

Berger, C., Renner, S., Luer, K., and Technau, G.M. (2007). The commonly used marker ELAV is transiently expressed in neuroblasts and glial cells in the Drosophila embryonic CNS. Dev Dyn 236, 3562–3568.

Bolte, S., and Cordelieres, F.P. (2006). A guided tour into subcellular colocalization analysis in light microscopy. J Microsc 224, 213–232.

Bornelov, S., Reynolds, N., Xenophontos, M., Gharbi, S., Johnstone, E., Floyd, R., Ralser, M., Signolet, J., Loos, R., Dietmann, S., et al. (2018). The Nucleosome Remodeling and Deacetylation Complex Modulates Chromatin Structure at Sites of Active Transcription to Fine-Tune Gene Expression. Mol Cell 71, 56–72 e54.

Brehm, A., Langst, G., Kehle, J., Clapier, C.R., Imhof, A., Eberharter, A., Muller, J., and Becker, P.B. (2000). dMi-2 and ISWI chromatin remodelling factors have distinct nucleosome binding and mobilization properties. EMBO J 19, 4332–4341.

Burgold, T., Barber, M., Kloet, S., Cramard, J., Gharbi, S., Floyd, R., Kinoshita, M., Ralser, M., Vermeulen, M., Reynolds, N., et al. (2019). The Nucleosome Remodelling and Deacetylation complex suppresses transcriptional noise during lineage commitment. EMBO J 38.

Dobin, A., Davis, C.A., Schlesinger, F., Drenkow, J., Zaleski, C., Jha, S., Batut, P., Chaisson, M., and Gingeras, T.R. (2013). STAR: ultrafast universal RNA-seq aligner. Bioinformatics 29, 15–21.

Eichler, K., Li, F., Litwin-Kumar, A., Park, Y., Andrade, I., Schneider-Mizell, C.M., Saumweber, T., Huser, A., Eschbach, C., Gerber, B., et al. (2017). The complete connectome of a learning and memory centre in an insect brain. Nature 548, 175–182.

Estacio-Gomez, A., Hassan, A., Walmsley, E., Le, L.W., and Southall, T.D. (2020). Dynamic neurotransmitter specific transcription factor expression profiles during Drosophila development. Biol Open 9.

Favorov, A., Mularoni, L., Cope, L.M., Medvedeva, Y., Mironov, A.A., Makeev, V.J., and Wheelan, S.J. (2012). Exploring massive, genome scale datasets with the GenometriCorr package. PLoS Comput Biol 8, e1002529.

Filion, G.J., van Bemmel, J.G., Braunschweig, U., Talhout, W., Kind, J., Ward, L.D., Brugman, W., de Castro, I.J., Kerkhoven, R.M., Bussemaker, H.J., et al. (2010). Systematic protein location mapping reveals five principal chromatin types in Drosophila cells. Cell 143, 212–224.

Gomez-Del Arco, P., Perdiguero, E., Yunes-Leites, P.S., Acin-Perez, R., Zeini, M., Garcia-Gomez, A., Sreenivasan, K., Jimenez-Alcazar, M., Segales, J., Lopez-Maderuelo, D., et al. (2016). The Chromatin Remodeling Complex Chd4/NuRD Controls Striated Muscle Identity and Metabolic Homeostasis. Cell Metab 23, 881–892.

Hassan, A., Araguas Rodriguez, P., Heidmann, S.K., Walmsley, E.L., Aughey, G.N., and Southall, T.D. (2020). Condensin I subunit Cap-G is essential for proper gene expression during the maturation of post-mitotic neurons. Elife 9.

Herrmann, C., Van de Sande, B., Potier, D., and Aerts, S. (2012). i-cisTarget: an integrative genomics method for the prediction of regulatory features and cis-regulatory modules. Nucleic Acids Res 40, e114.

Ho, J.W., Jung, Y.L., Liu, T., Alver, B.H., Lee, S., Ikegami, K., Sohn, K.A., Minoda, A., Tolstorukov, M.Y., Appert, A., et al. (2014). Comparative analysis of metazoan chromatin organization. Nature 512, 449–452.

Hoffmann, A., and Spengler, D. (2019). Chromatin Remodeling Complex NuRD in Neurodevelopment and Neurodevelopmental Disorders. Front Genet 10, 682.

Hsu, T., Bagni, C., Sutherland, J.D., and Kafatos, F.C. (1996). The transcriptional factor CF2 is a mediator of EGF-R-activated dorsoventral patterning in Drosophila oogenesis. Genes Dev 10, 1411–1421.

Hu, Y., Comjean, A., Perrimon, N., and Mohr, S.E. (2017). The Drosophila Gene Expression Tool (DGET) for expression analyses. BMC Bioinformatics 18, 98.

Huang, Z., and Kunes, S. (1996). Hedgehog, transmitted along retinal axons, triggers neurogenesis in the developing visual centers of the Drosophila brain. Cell 86, 411–422.

Janssens, J., Aibar, S., Taskiran, II, Ismail, J.N., Gomez, A.E., Aughey, G., Spanier, K.I., De Rop, F.V., Gonzalez-Blas, C.B., Dionne, M., et al. (2022). Decoding gene regulation in the fly brain. Nature.

Jiao, W., Spreemann, G., Ruchti, E., Banerjee, S., Shi, Y., Stowers, R.S., Hess, K., and McCabe, B.D. (2021). Intact Drosophila Whole Brain Cellular Quantitation reveals Sexual Dimorphism. bioRxiv, 2021.2011.2003.467146.

Johnson, P.W., Doe, C.Q., and Lai, S.L. (2018). Drosophila nucleostemin 3 is required to maintain larval neuroblast proliferation. Dev Biol 440, 1–12.

Krause, S.A., Overend, G., Dow, J.A.T., and Leader, D.P. (2022). FlyAtlas 2 in 2022: enhancements to the Drosophila melanogaster expression atlas. Nucleic Acids Res 50, D1010–D1015.

Kreher, J., Kovac, K., Bouazoune, K., Macinkovic, I., Ernst, A.L., Engelen, E., Pahl, R., Finkernagel, F., Murawska, M., Ullah, I., et al. (2017). EcR recruits dMi-2 and increases efficiency of dMi-2-mediated remodelling to constrain transcription of hormone-regulated genes. Nat Commun 8, 14806.

Kunert, N., Wagner, E., Murawska, M., Klinker, H., Kremmer, E., and Brehm, A. (2009). dMec: a novel Mi-2 chromatin remodelling complex involved in transcriptional repression. EMBO J 28, 533–544.

Lattao, R., Kovacs, L., and Glover, D.M. (2017). The Centrioles, Centrosomes, Basal Bodies, and Cilia of Drosophila melanogaster. Genetics 206, 33–53.

Liao, Y., Smyth, G.K., and Shi, W. (2019). The R package Rsubread is easier, faster, cheaper and better for alignment and quantification of RNA sequencing reads. Nucleic Acids Res 47, e47.

Love, M.I., Huber, W., and Anders, S. (2014). Moderated estimation of fold change and dispersion for RNA-seq data with DESeq2. Genome Biol 15, 550.

Low, J.K., Webb, S.R., Silva, A.P., Saathoff, H., Ryan, D.P., Torrado, M., Brofelth, M., Parker, B.L., Shepherd, N.E., and Mackay, J.P. (2016). CHD4 Is a Peripheral Component of the Nucleosome Remodeling and Deacetylase Complex. J Biol Chem 291, 15853–15866.

Marshall, O.J., and Brand, A.H. (2015). damidseq_pipeline: an automated pipeline for processing DamID sequencing datasets. Bioinformatics 31, 3371–3373.

Marshall, O.J., and Brand, A.H. (2017). Chromatin state changes during neural development revealed by in vivo cell-type specific profiling. Nat Commun 8, 2271.

Marshall, O.J., Southall, T.D., Cheetham, S.W., and Brand, A.H. (2016). Cell-type-specific profiling of protein-DNA interactions without cell isolation using targeted DamID with next-generation sequencing. Nat Protoc 11, 1586–1598.

Nichols, C.D., Becnel, J., and Pandey, U.B. (2012). Methods to assay Drosophila behavior. J Vis Exp.

Nitarska, J., Smith, J.G., Sherlock, W.T., Hillege, M.M., Nott, A., Barshop, W.D., Vashisht, A.A., Wohlschlegel, J.A., Mitter, R., and Riccio, A. (2016). A Functional Switch of NuRD Chromatin Remodeling Complex Subunits Regulates Mouse Cortical Development. Cell Rep 17, 1683–1698.

O’Shaughnessy-Kirwan, A., Signolet, J., Costello, I., Gharbi, S., and Hendrich, B. (2015). Constraint of gene expression by the chromatin remodelling protein CHD4 facilitates lineage specification. Development 142, 2586–2597.

Preissl, S., Fang, R., Huang, H., Zhao, Y., Raviram, R., Gorkin, D.U., Zhang, Y., Sos, B.C., Afzal, V., Dickel, D.E., et al. (2018). Single-nucleus analysis of accessible chromatin in developing mouse forebrain reveals cell-type-specific transcriptional regulation. Nat Neurosci 21, 432–439.

Quinlan, A.R., and Hall, I.M. (2010). BEDTools: a flexible suite of utilities for comparing genomic features. Bioinformatics 26, 841–842.

Ragheb, R., Gharbi, S., Cramard, J., Ogundele, O., Kloet, S.L., Burgold, T., Vermeulen, M., Reynolds, N., and Hendrich, B. (2020). Differential regulation of lineage commitment in human and mouse primed pluripotent stem cells by the nucleosome remodelling and deacetylation complex. Stem Cell Res 46, 101867.

Rais, Y., Zviran, A., Geula, S., Gafni, O., Chomsky, E., Viukov, S., Mansour, A.A., Caspi, I., Krupalnik, V., Zerbib, M., et al. (2013). Deterministic direct reprogramming of somatic cells to pluripotency. Nature 502, 65–70.

Ravenscroft, T.A., Janssens, J., Lee, P.T., Tepe, B., Marcogliese, P.C., Makhzami, S., Holmes, T.C., Aerts, S., and Bellen, H.J. (2020). Drosophila Voltage-Gated Sodium Channels Are Only Expressed in Active Neurons and Are Localized to Distal Axonal Initial Segment-like Domains. J Neurosci 40, 7999–8024.

Reddington, J.P., Garfield, D.A., Sigalova, O.M., Karabacak Calviello, A., Marco-Ferreres, R., Girardot, C., Viales, R.R., Degner, J.F., Ohler, U., and Furlong, E.E.M. (2020). Lineage-Resolved Enhancer and Promoter Usage during a Time Course of Embryogenesis. Dev Cell 55, 648–664 e649.

Reddy, B.A., Bajpe, P.K., Bassett, A., Moshkin, Y.M., Kozhevnikova, E., Bezstarosti, K., Demmers, J.A., Travers, A.A., and Verrijzer, C.P. (2010). Drosophila transcription factor Tramtrack69 binds MEP1 to recruit the chromatin remodeler NuRD. Mol Cell Biol 30, 5234–5244.

Rhee, D.Y., Cho, D.Y., Zhai, B., Slattery, M., Ma, L., Mintseris, J., Wong, C.Y., White, K.P., Celniker, S.E., Przytycka, T.M., et al. (2014). Transcription factor networks in Drosophila melanogaster. Cell Rep 8, 2031–2043.

Selleck, S.B., and Steller, H. (1991). The influence of retinal innervation on neurogenesis in the first optic ganglion of Drosophila. Neuron 6, 83–99.

Southall, T.D., Davidson, C.M., Miller, C., Carr, A., and Brand, A.H. (2014). Dedifferentiation of neurons precedes tumor formation in Lola mutants. Dev Cell 28, 685–696.

Southall, T.D., Gold, K.S., Egger, B., Davidson, C.M., Caygill, E.E., Marshall, O.J., and Brand, A.H. (2013). Cell-type-specific profiling of gene expression and chromatin binding without cell isolation: assaying RNA Pol II occupancy in neural stem cells. Dev Cell 26, 101–112.

Tie, F., Furuyama, T., Prasad-Sinha, J., Jane, E., and Harte, P.J. (2001). The Drosophila Polycomb Group proteins ESC and E(Z) are present in a complex containing the histone-binding protein p55 and the histone deacetylase RPD3. Development 128, 275–286.

Tong, J.K., Hassig, C.A., Schnitzler, G.R., Kingston, R.E., and Schreiber, S.L. (1998). Chromatin deacetylation by an ATP-dependent nucleosome remodelling complex. Nature 395, 917–921.

Unhavaithaya, Y., Shin, T.H., Miliaras, N., Lee, J., Oyama, T., and Mello, C.C. (2002). MEP-1 and a homolog of the NURD complex component Mi-2 act together to maintain germline-soma distinctions in C. elegans. Cell 111, 991–1002.

Wade, P.A., Jones, P.L., Vermaak, D., and Wolffe, A.P. (1998). A multiple subunit Mi-2 histone deacetylase from Xenopus laevis cofractionates with an associated Snf2 superfamily ATPase. Curr Biol 8, 843–846.

Wang, H.B., and Zhang, Y. (2001). Mi2, an auto-antigen for dermatomyositis, is an ATP-dependent nucleosome remodeling factor. Nucleic Acids Res 29, 2517–2521.

Wang, Z., Zang, C., Cui, K., Schones, D.E., Barski, A., Peng, W., and Zhao, K. (2009). Genome-wide mapping of HATs and HDACs reveals distinct functions in active and inactive genes. Cell 138, 1019–1031.

Xue, Y., Wong, J., Moreno, G.T., Young, M.K., Cote, J., and Wang, W. (1998). NURD, a novel complex with both ATP-dependent chromatin-remodeling and histone deacetylase activities. Mol Cell 2, 851–861.

Yamada, T., Yang, Y., Hemberg, M., Yoshida, T., Cho, H.Y., Murphy, J.P., Fioravante, D., Regehr, W.G., Gygi, S. P., Georgopoulos, K., et al. (2014). Promoter decommissioning by the NuRD chromatin remodeling complex triggers synaptic connectivity in the mammalian brain. Neuron 83, 122–134.

Yoshida, T., Hazan, I., Zhang, J., Ng, S.Y., Naito, T., Snippert, H.J., Heller, E.J., Qi, X., Lawton, L.N., Williams, C.J., et al. (2008). The role of the chromatin remodeler Mi-2beta in hematopoietic stem cell self-renewal and multilineage differentiation. Genes Dev 22, 1174–1189.

Zacharioudaki, E., Falo Sanjuan, J., and Bray, S. (2019). Mi-2/NuRD complex protects stem cell progeny from mitogenic Notch signaling. Elife 8.

Zhang, W., Aubert, A., Gomez de Segura, J.M., Karuppasamy, M., Basu, S., Murthy, A.S., Diamante, A., Drury, T. A., Balmer, J., Cramard, J., et al. (2016). The Nucleosome Remodeling and Deacetylase Complex NuRD Is Built from Preformed Catalytically Active Sub-modules. J Mol Biol 428, 2931–2942.

Zhang, Y., LeRoy, G., Seelig, H.P., Lane, W.S., and Reinberg, D. (1998). The dermatomyositis-specific autoantigen Mi2 is a component of a complex containing histone deacetylase and nucleosome remodeling activities. Cell 95, 279–289.

Ziffra, R.S., Kim, C.N., Ross, J.M., Wilfert, A., Turner, T.N., Haeussler, M., Casella, A.M., Przytycki, P.F., Keough, K.C., Shin, D., et al. (2021). Single-cell epigenomics reveals mechanisms of human cortical development. Nature 598, 205–213.

