## Supplementary material for "NuRD independent Mi-2 activity represses ectopic gene expression during neuronal maturation": Expanded view figures

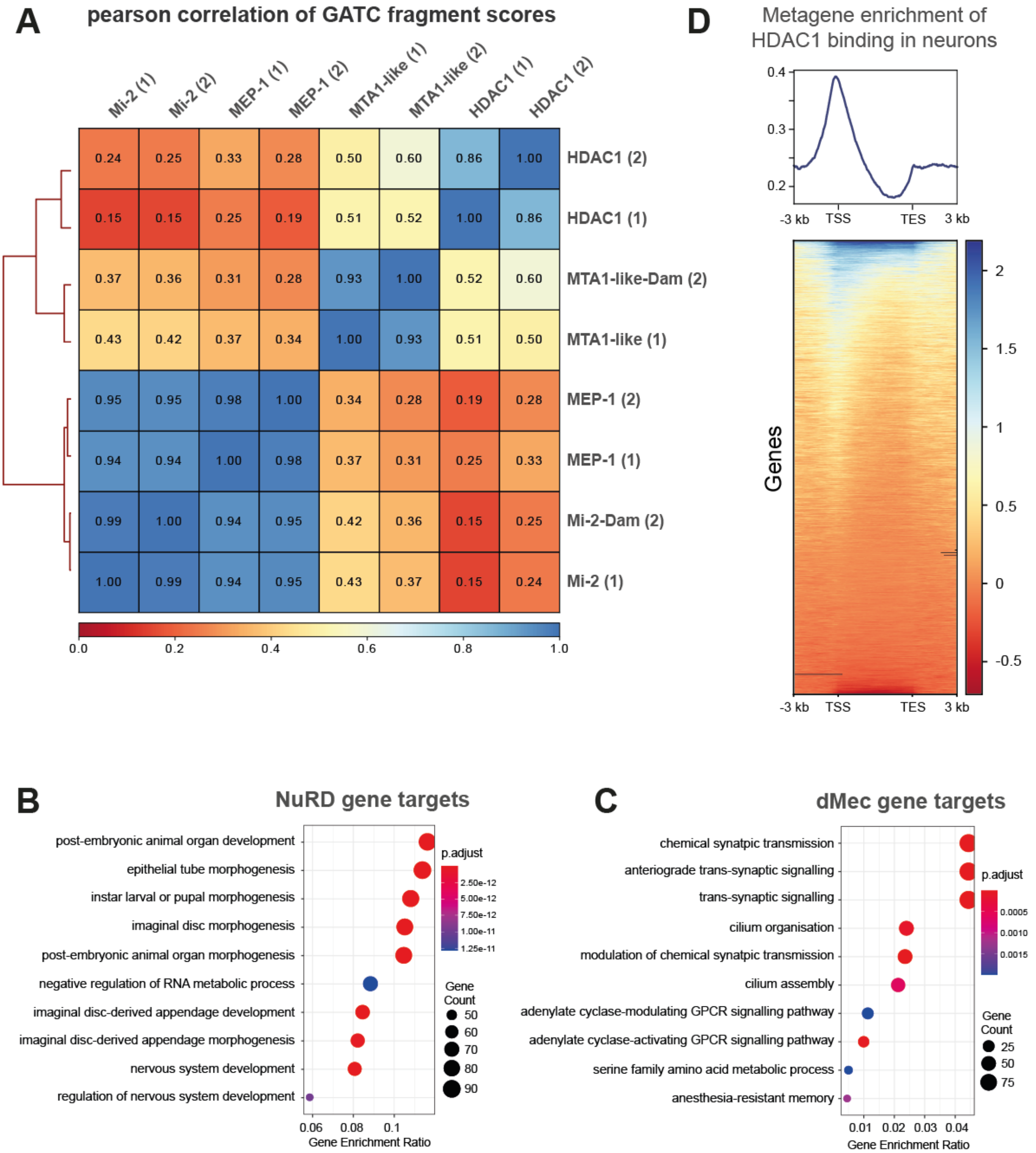


**Figure EV1. Genomic binding of NuRD components.** A) Correlation between NuRD subunits (including replicates) in larval neurons (*elav*-GAL4). Very strong correlations are seen between Mi-2 and MEP-1 binding while the correlation between Mi-2/MEP-1 and either MTA-1like or HDAC1 are relatively lower. Good correlations are exhibited between replicates for each subunit (spearman's correlation $r^2$=0.85-0.95). B) GO analysis of NuRD gene targets. C) GO analysis of dMec gene targets genes. D) Metagene analysis of HDAC1 binding. TSS is Transcriptional Start Site and TES is Transcriptional End Site.

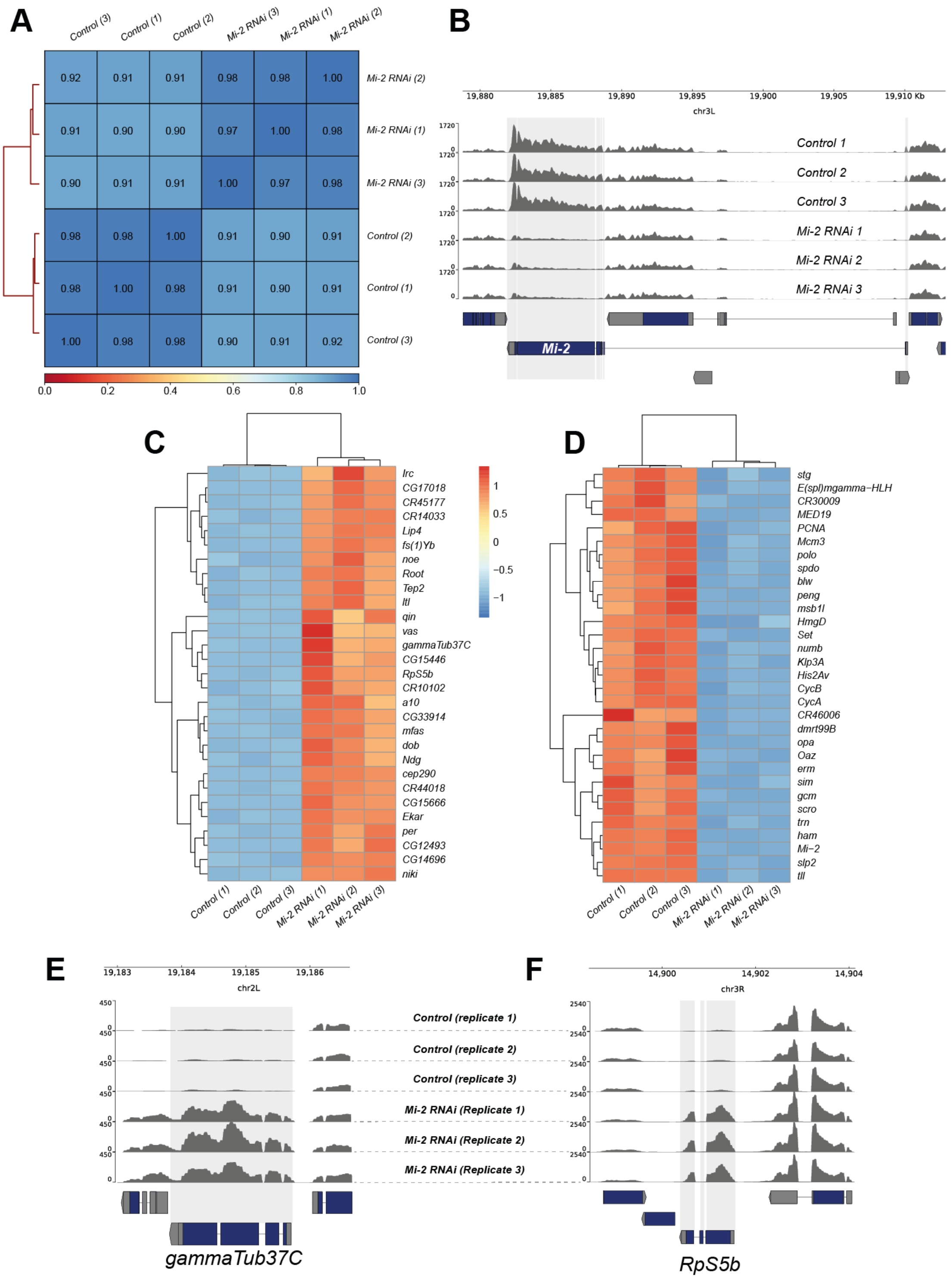


**Figure EV2. RNA-seq quality control and differentially expressed genes.** A) Heatmap showing Pearson correlation of RNA-seq reads between biological replicates and control and experimental groups. Very strong correlations are observed between replicates ($R^2 > 0.97$) with slightly lower correlations between knockdown and controls reflecting the changes in *Mi-2* RNAi transcriptome. B) Genome browser tracks indicating RNA-seq read coverage at the *Mi-2* locus. *Mi-2* reads are clearly depleted in *Mi-2 RNAi* compared to *elav*-GAL4 *x mCherry RNAi* controls (adjusted p value < 2.2 x $10^{-227}$). C) Heatmap showing relative changes in normalised gene expression for the thirty most significantly up-regulated genes. D) Heatmap showing relative changes in normalised gene expression for the thirty most significantly down-regulated genes. E) Read coverage for E) *gammaTub37C* and F) *RpS5b* loci, which are non-CNS genes ectopically expressed in *Mi-2* knockdown.

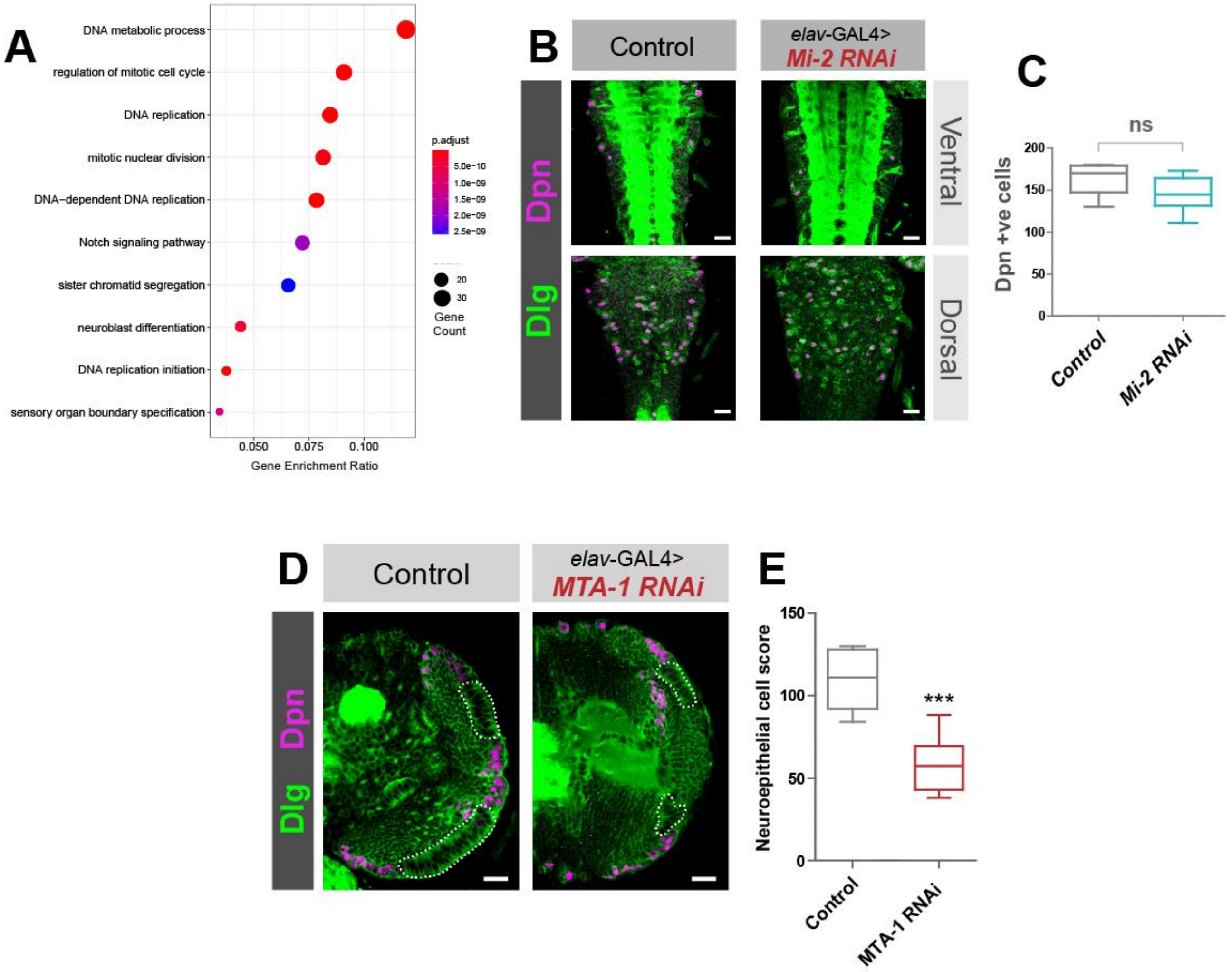


**Figure EV3. *NuRD* knockdown larval brain phenotypes.** A) GO analysis of down-regulated genes. B) *Mi-2* knockdown does not affect NSC numbers in the VNC. B) Example images from ventral and dorsal sections of the VNC. Scale bars = 20 µm. C) Quantification of Dpn positive cells (no significant difference). 6 VNCs measured for each genotype. No significant difference (two-tailed student's t-test). Represented as a box plot. D) *MTA1-like* knockdown causes an optic lobe phenotype. D) Representative images of control (*elav*-GAL4; ; *mCherry RNAi*), and *MTA1-like* knockdown in the third instar larval CNS (all scale bars = 20 µm). Discs large (Dlg) = green, Deadpan (Dpn) = magenta. Neuroepithelial cells are highlighted by dashed lines. E) Quantification of optic lobe neuroepithelial cells of genotypes shown in panel A. At least 8 brains measured for each genotype. *** $p < 0.001$ (one-tailed student's t-test). Represented as a box plot.

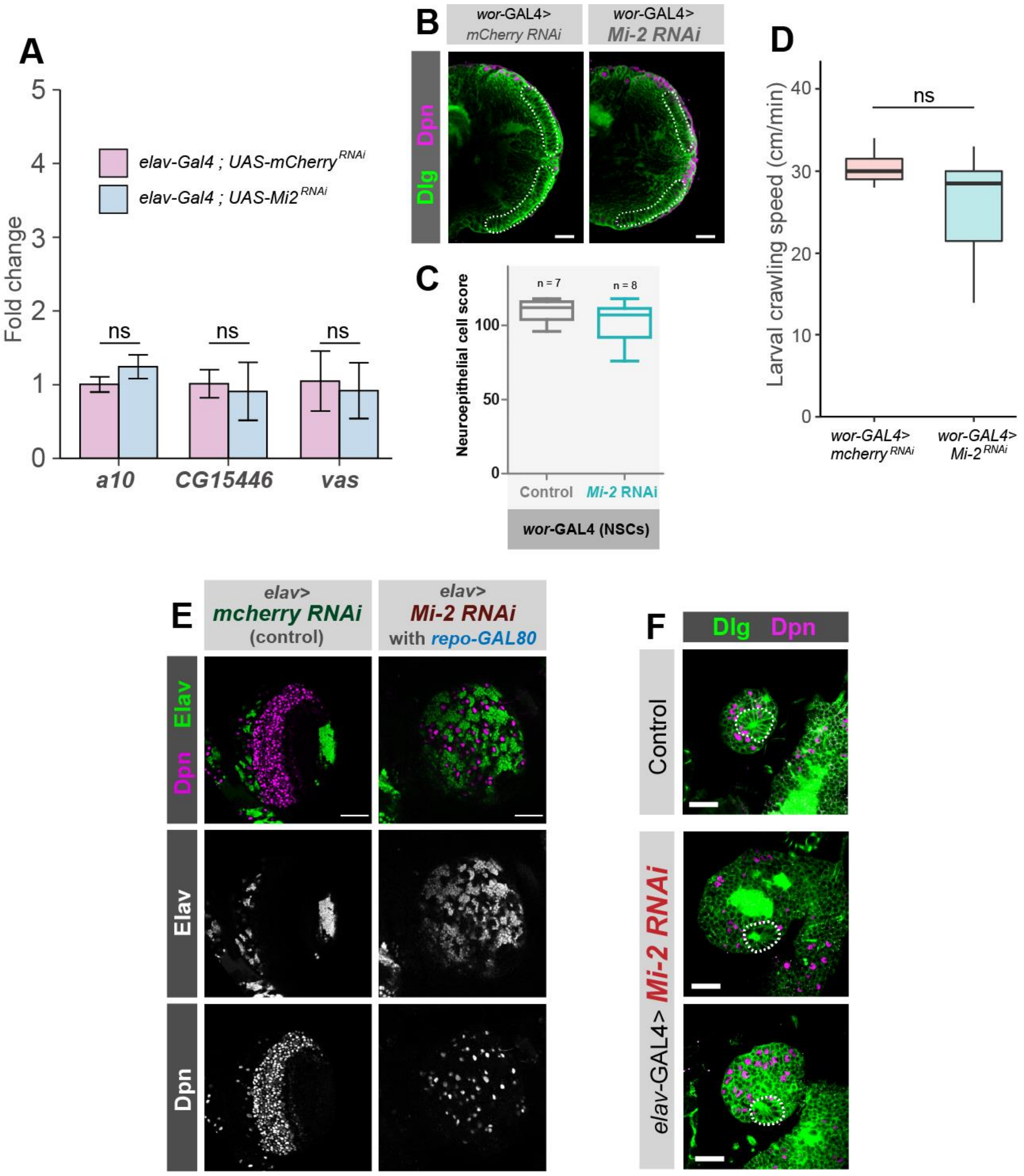


**Figure EV4. Differential requirements of Mi-2 during neuronal differentiation.** A) Knockdown of *Mi-2* in adult brain neurons does not cause ectopic expression of genes. qPCR measurement of gene expression in adult heads. *Mi-2*$^{RNAi}$ is induced for 24 hours using GAL80$^{ts}$. No significant changes in expression of either *vas, a10,* or *CG15446* were observed (one-tailed students t-test, represented as mean ± SEM.). 3 biological replicates per genotype. B) Representative images of optic lobes for control and *Mi-2* knockdown in NSCs. C) Quantification of optic lobe neuroepithelial cells of genotypes shown in panel B (n >= 7 for each group). Represented as a box plot. No significant difference found (one-tailed student's t-test). D) Larval crawl speed (cm/minute) in *wor*-GAL4 x *Mi-2* RNAi and *wor*-GAL4 x *mCherry* RNAi controls. No significant changes in crawl speed were observed (two-tailed students t-test, n = 10 animals per genotype). Represented as a box plot. E) Optic lobe phenotype is not due to *Mi-2* knockdown in glial cells. Optic lobe of third instar larvae in which *repo*-GAL80 represses glial GAL4 expression. Loss of neuroepithelial cells is still observed as in *elav*-GAL4 *; Mi-2 RNAi* brains. Scale bar = 50 µm. F) Characteristic rosette structure of the neuroepithelial cells are present in the control 1$^{st}$ instar optic lobe and in *elav*-GAL4 *; Mi-2 RNAi* 1$^{st}$ instar optic lobes. Scale bar = 20 µm.

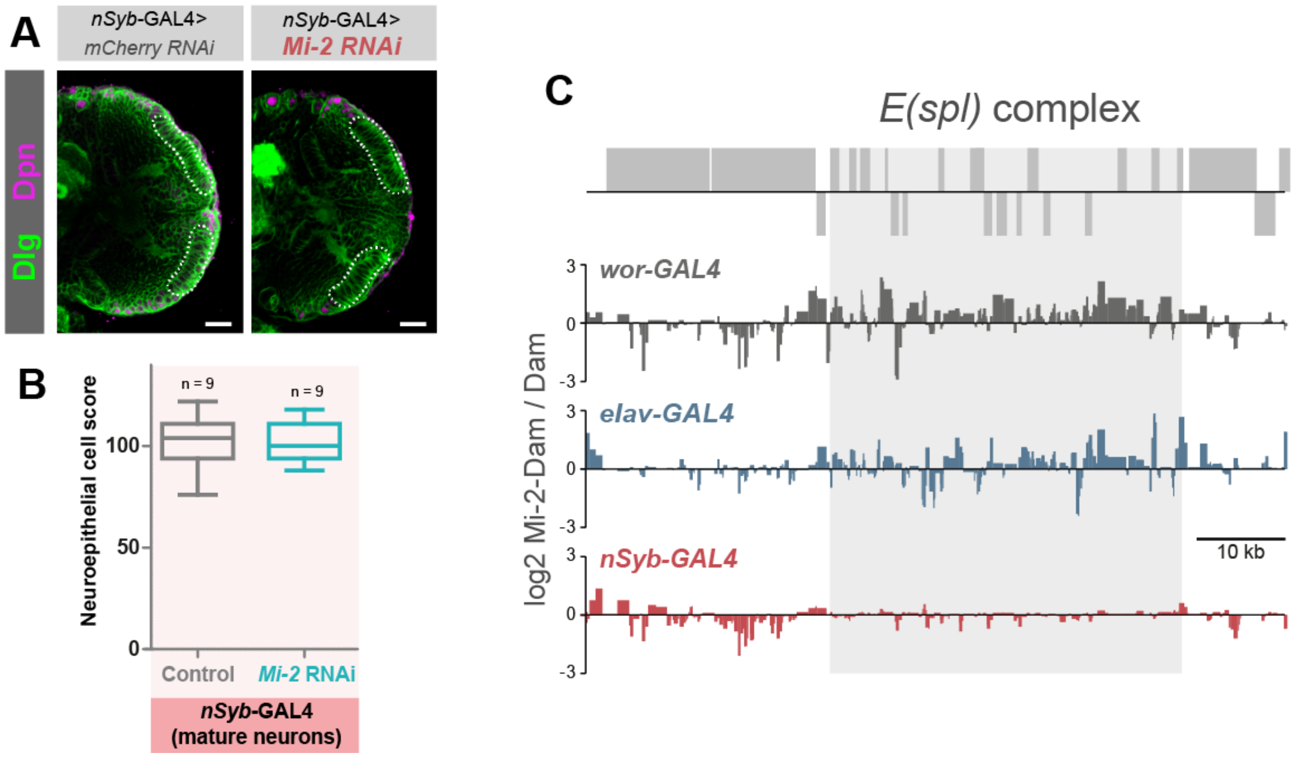


**Figure EV5. Differential requirements and binding of Mi-2 during neuronal differentiation.** A) Representative images of optic lobes for control and *Mi-2* knockdown in mature neurons. B) Quantification of optic lobe neuroepithelial cells of genotypes shown in panel B (n >= 7 for each group). Represented as a box plot. No significant difference found (one-tailed student's t-test). C) Mi-2 binding at the *E(spl)* complex is lost in mature neurons. Grey boxes represent genes.
