## Appendix Figures and Tables for "NuRD independent Mi-2 activity represses ectopic gene expression during neuronal maturation"

### Appendix information

**Contents:**

**Appendix Figure S1.** Tissue enrichment analysis for differentially upregulated genes associated with the “cilium organisation” GO term.

**Appendix Figure S2.** Expression of differentially expressed genes in other tissues.

**Appendix Figure S3.** Verification of *Mi-2* and *HDAC1* RNAi lines.

**Appendix Figure S4.** Pupal lethality in NuRD component RNAi knockdown experiments

**Appendix Table S1.** Fly stocks used.

**Appendix Table S2.** DNA oligos used in this study.

**Appendix References**

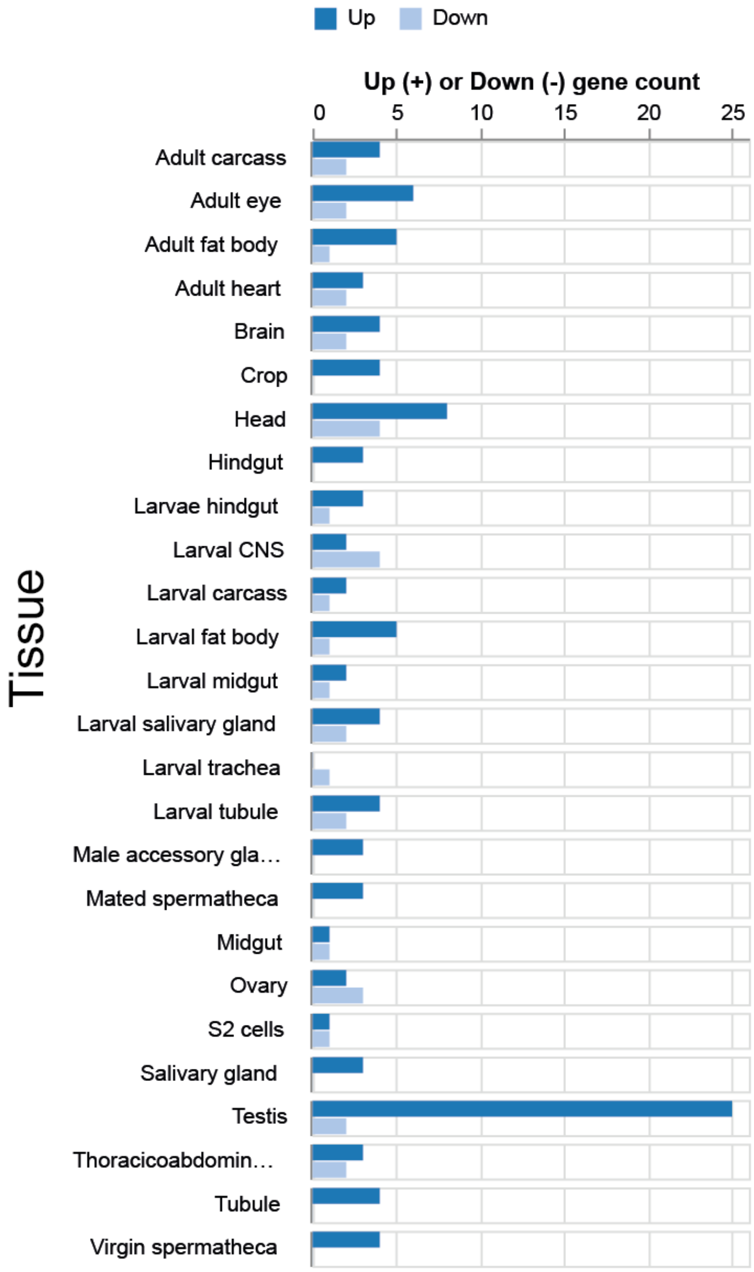


**Appendix figure S1. Tissue enrichment analysis for differentially upregulated genes associated with the "cilium organisation" GO term.** The majority of these genes are highly upregulated in testis. "Up" and "Down" refer to genes in the set that are significantly up or downregulated in the respective tissue.

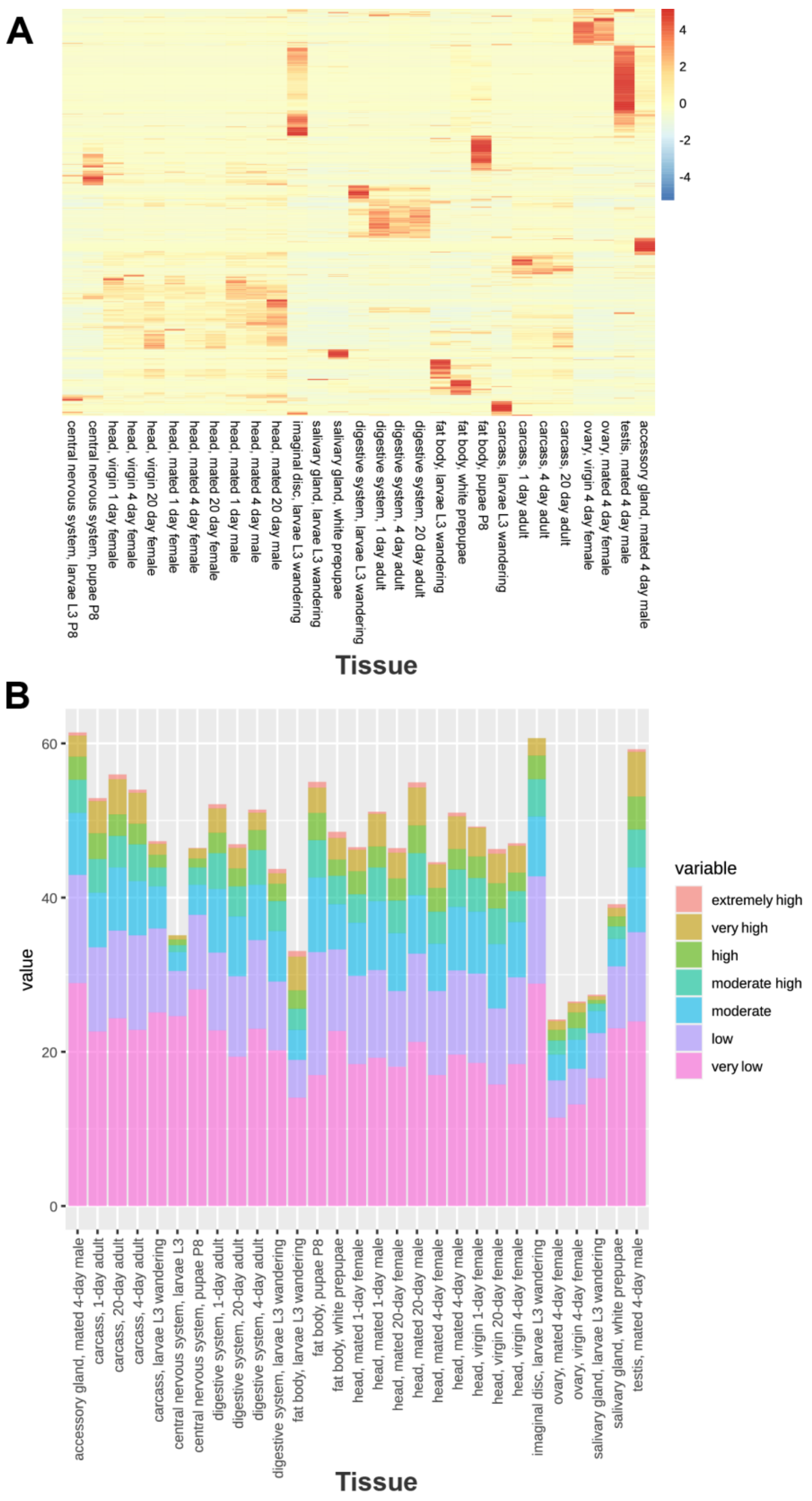


**Appendix figure S2. Expression of differentially expressed genes in other tissues.** A) Heatmap showing relative expression levels of upregulated genes across all tissues. B) Stacked bar plot showing percentage of upregulated genes that are expressed at different levels in specific tissues. Note that many upregulated genes are expressed at "very low" levels in the fly CNS, whilst upregulated genes are frequently expressed at "moderate" to very high levels in germline and other non-CNS tissues.

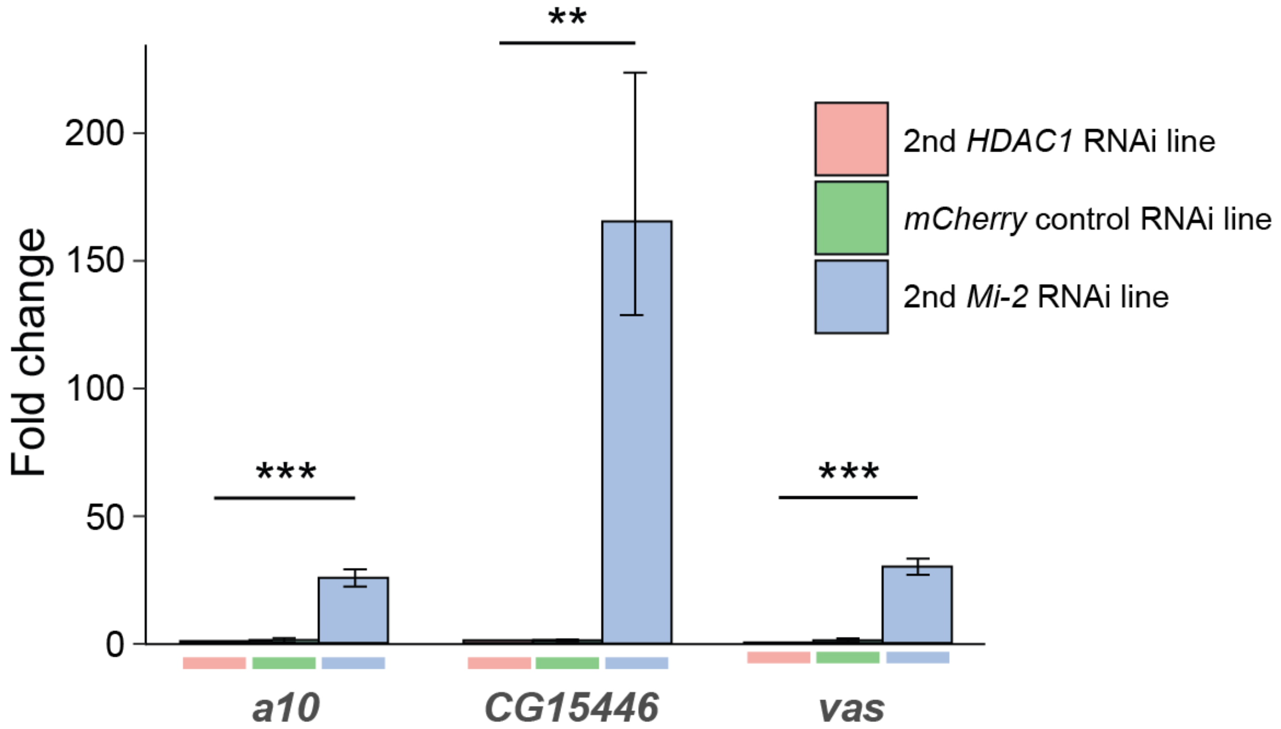


**Appendix figure S3. Verification of *Mi-2* and *HDAC1* RNAi lines.** qPCR measurement of non-neuronal genes in larval brains using alternative RNAi lines for *Mi-2* and *HDAC* (n = 3). These confirm the results obtained using the original RNAi lines. **P < 0.005, ***P < 0.0005 (one-tailed student's t-test). Error bars show standard deviation. Represented as mean ± SEM.

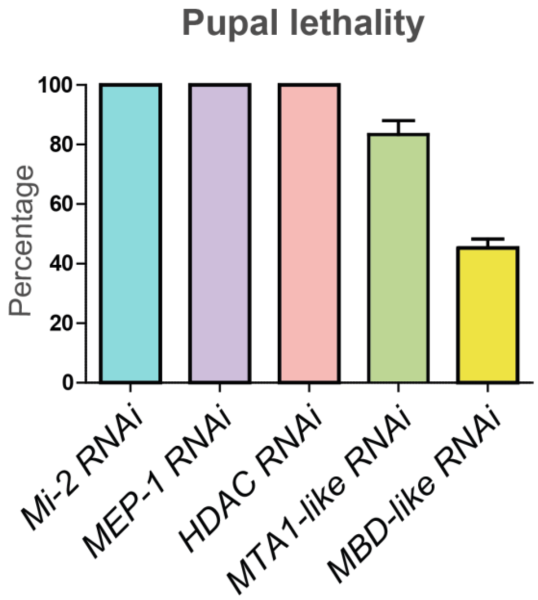


**Appendix figure S4. Pupal lethality in NuRD component RNAi knockdown experiments.** Percentage of pupae that die when NuRD subunits are knocked down in neurons (driven by *elav*-GAL4) during development. 3 independent experiments were performed for each genotype.

**Appendix Table S1. Fly stocks used.**

| Genotype | Source | Reference | Description |
|---|---|---|---|
| *$y^1w^*$; Mi{PT-GFSTF.1}Mi-2$^{MI07934\text{-}GFSTF.1}$ Su(Tpl)$^{MI07934\text{-}GFSTF.1\text{-}X}$/TM6C, Sb$^1$ Tb$^1$* | BDSC #63188 | (Nagarkar-Jaiswal et al., 2015) | Mi-2 GFP trap |
| *$y^1$ $w^*$; Mi{PT-GFSTF.1}MTA1-like$^{MI01790\text{-}GFSTF.1}$* | BDSC #63161 | (Nagarkar-Jaiswal et al., 2015) | MTA GFP trap |
| *UAS-LT3*-Dam | Andrea Brand | (Southall et al., 2013) | |
| *UAS*- Dam-MEP-1 | This study | | |
| *UAS*-Dam-Mi-2 | This study | | |
| *UAS*-MTA1-like-Dam | This study | | flyORF TaDa |
| *UAS*-HDAC-Dam | This study | | flyORF TaDa |
| *wor*-GAL4; *tub*-GAL80$^{ts}$ | Andrea Brand | (Albertson et al., 2004) | |
| *wor*-GAL4 | Andrea Brand | (Albertson et al., 2004) | |
| *elav*-GAL4*; tub*-GAL80$^{ts}$ | Andrea Brand | | |
| *nSyb*-GAL4 | BDSC #51941 | | |
| *hs-flp; UAS-flyORF.TaDa* | BDSC #91637 | (Aughey et al., 2021) | |
| *UAS*-HDAC1 | FlyORF F000675 | (Bischof et al., 2013) | |
| *UAS*-MTA1-like | FlyORF F001892 | (Bischof et al., 2013) | |
| *UAS-Mi2 RNAi* | BDSC #51774 | | |
| *UAS-Mi2 RNAi* (2nd line) | BDSC #35398 | | |
| *UAS-MTA1-like RNAi* | BDSC #34624 | | |
| *UAS-HDAC1 RNAi* | BDSC #33725 | | |
| *UAS-HDAC1 RNAi* (2nd line) | BDSC #34846 | | |
| *UAS-MEP-1 RNAi* | BDSC #62180 | (Perkins et al., 2015) | |
| *UAS-MBD-like RNAi* | VDRC #9261 | (Dietzl et al., 2007) | |
| *mCherry-RNAi (control)* | BDSC #35787 | | |
| *repo*-GAL80 | Manolis Fanto | (Awasaki et al., 2008) | |
| *elav*-GAL4 ; *repo*-GAL80 / *TM6* | This study | | |

**Appendix Table S2. DNA oligos used in this study.**

| **Primer name** | **Primer sequence** |
| --- | --- |
| *vas* qRT-PCR forward | TGTCTGACGACTGGGATGATG |
| *vas* qRT-PCR reverse | ATTTCCTCCTTGGTAGCCGC |
| *CG17566* qRT-PCR forward | TTGGGCCAATGTGGCAATCA |
| *CG17566* qRT-PCR reverse | TGCGATCCTGTCCATCGGT |
| *qin* qRT-PCR forward | TCCCTTTCTACTGGGATGCG |
| *qin* qRT-PCR reverse | GAGCTGGACTATGGCACACG |
| *RpS5b* qRT-PCR forward | ACTACATTGCCGTAAAGGAGAAG |
| *RpS5b* qRT-PCR reverse | CATTGGGCCTTGCGGAATC |
| *CG8526* qRT-PCR forward | GATTTGATTGAGTTCTGCCCACT |
| *CG8526* qRT-PCR reverse | CTTGTATGTCTTTCCCACTTCGT |
| *CG15446* qRT-PCR forward | CGGAAACGGCTACCCATGT |
| *CG15446* qRT-PCR reverse | CCCCGACTTACCTTCATCTTCG |
| *a10* qRT-PCR forward | ATCCTTAACCAAGAGCGACTGT |
| *a10* qRT-PCR reverse | TCACCTTTTCAGCACCATACC |
| *RpL4* qRT-PCR forward | TCCACCTTGAAGAAGGGCTA |
| *RpL4* qRT-PCR reverse | TTGCGGATCTCCTCAGACTT |
| *Dam-Mi2*_forward | gaagaggatctggccggcgcagatctgcggATGGCATCGGAGGAAGAGAATGACGATAAT |
| *Dam-Mi2*_reverse | aagtaaggttccttcacaaagatcctctagCTAGACGCCGGAATTATTCGATAGCTGGCC |
| *Dam-MEP-1*_forward | gaagaggatctggccggcgcagatctgcggATGACTGAAGTTGATGTCGTTTTGCCGGAG |
| *Dam-MEP-1*_reverse | aagtaaggttccttcacaaagatcctctagTTAATCTATGACATGACTCTCCATATTTG |

### Appendix References


Albertson, R., Chabu, C., Sheehan, A., and Doe, C.Q. (2004). Scribble protein domain mapping reveals a multistep localization mechanism and domains necessary for establishing cortical polarity. *J Cell Sci* 117, 6061-6070.

Aughey, G.N., Delandre, C., McMullen, J.P.D., Southall, T.D., and Marshall, O.J. (2021). FlyORF-TaDa allows rapid generation of new lines for in vivo cell-type-specific profiling of protein-DNA interactions in Drosophila melanogaster. *G3 (Bethesda)* 11.

Awasaki, T., Lai, S.L., Ito, K., and Lee, T. (2008). Organization and postembryonic development of glial cells in the adult central brain of Drosophila. *J Neurosci* 28, 13742-13753.

Bischof, J., Bjorklund, M., Furger, E., Schertel, C., Taipale, J., and Basler, K. (2013). A versatile platform for creating a comprehensive UAS-ORFeome library in Drosophila. *Development* 140, 2434-2442.

Dietzl, G., Chen, D., Schnorrer, F., Su, K.C., Barinova, Y., Fellner, M., Gasser, B., Kinsey, K., Oppel, S., Scheiblauer, S.*, et al.* (2007). A genome-wide transgenic RNAi library for conditional gene inactivation in Drosophila. *Nature* 448, 151-156.

Nagarkar-Jaiswal, S., Lee, P.T., Campbell, M.E., Chen, K., Anguiano-Zarate, S., Gutierrez, M.C., Busby, T., Lin, W.W., He, Y., Schulze, K.L.*, et al.* (2015). A library of MiMICs allows tagging of genes and reversible, spatial and temporal knockdown of proteins in Drosophila. *Elife* 4.

Perkins, L.A., Holderbaum, L., Tao, R., Hu, Y., Sopko, R., McCall, K., Yang-Zhou, D., Flockhart, I., Binari, R., Shim, H.S.*, et al.* (2015). The Transgenic RNAi Project at Harvard Medical School: Resources and Validation. *Genetics* 201, 843-852.

Southall, T.D., Gold, K.S., Egger, B., Davidson, C.M., Caygill, E.E., Marshall, O.J., and Brand, A.H. (2013). Cell-type-specific profiling of gene expression and chromatin binding without cell isolation: assaying RNA Pol II occupancy in neural stem cells. *Dev Cell* 26, 101-112.
